# Emergent population activity in metric-free and metric networks of neurons with stochastic spontaneous spikes and dynamic synapses

**DOI:** 10.1101/2021.05.08.442778

**Authors:** Dmitrii Zendrikov, Alexander Paraskevov

## Abstract

We show that networks of excitatory neurons with stochastic spontaneous spiking activity and short-term synaptic plasticity can exhibit spontaneous repetitive synchronization in so-called population spikes. The major reason for this is that synaptic plasticity nonlinearly modulates the interaction between neurons. For large-scale two-dimensional networks, where the connection probability decreases exponentially with increasing distance between the neurons resulting in a small-world network connectome, a population spike occurs in the form of circular traveling waves diverging from seemingly non-stationary nucleation sites. The latter is in drastic contrast to the case of networks with a fixed fraction of steady pacemaker neurons, where the set of a few spontaneously formed nucleation sites is stationary. Despite the spatial non-stationarity of their nucleation, population spikes may occur surprisingly regularly. From a theoretical viewpoint, these findings show that the regime of nearly-periodic population spikes, which mimics respiratory rhythm, can occur strictly without stochastic resonance. In addition, the observed spatiotemporal effects serve as an example of transient chimera patterns.

## 1. Introduction

It is now generally accepted that synaptic plasticity is a key property of biological neuronal networks, which determines their unsurpassed ability to process information [1]. For example, it has been assumed that short-term synaptic plasticity [2] could be the basis of working memory [3, 4]. On the other hand, abnormal manifestations of synaptic plasticity may lead to severe consequences, e.g., to episodic spontaneous pathological synchronization of spiking activity of neurons that underlies epilepsy [5].

Numerical simulations provide invaluable assistance in identifying the causes of emergent collective phenomena based on synaptic plasticity. One of the phenomena that we focus on in this paper is the regime of repetitive spontaneous population spikes (PSs) typical for planar neuronal networks cultured *in vitro* [6, 7]. Despite the fact that numerical simulation of the PS regime is easily accessible and there are numerous theoretical prerequisites for explaining this phenomenon (e.g., [8–19]), a clear self-consistent and predictive theory of the emergence of the PS regime is not yet available even in relatively simple neuronal network models. It is worth noting that many theories based on mean-field or rate-based models naturally lose the irregularity of PS occurrence (e.g., [20–22]) and, in the majority of cases, do not have the same statistical properties as full-featured spiking neuronal network models (cp. [23]).

In turn, spike-based models demonstrating this regime are usually deterministic [24–30], i.e. the dynamics of each neuron is described by deterministic equations (for a broader overview, see [31]). Given this, the ‘individuality’ of each specific neuron is determined either by additive noise to the synaptic current [32–35], or by an individual constant ‘background’ current, the value of which regulates the magnitude of neuronal excitability and the fraction of pacemaker neurons [24–26, 29, 36, 37]. In the case of background currents, the neuronal network model is entirely deterministic, yet it simulates the PS regime surprisingly well (and, importantly, completely reproducibly), including the formation of PS nucleation sites [29, 37].

However, experiments with brain slices indicate that there are two types of PSs in the brain: one is deterministic, as in neuronal cultures and probably in their deterministic models, and the other is stochastic [38]. (It is called stochastic because of the absence of any repeating characteristic pattern of spiking activity preceding the occurrence of a PS, cp. [39].) Also, there exists a disagreement between experimental findings even for cultured networks: PS spatial sources seemed steady for a given network of spinal cord neurons [40–42] while these were estimated as random for a network of cortical neurons [43]. A possible stochastic mechanism of synchronized cluster occurrence has been theoretically considered in Ref. [44] for a network of FitzHugh-Nagumo neurons with electrical synapses (i.e., gap junctions), phenomenological delay of interaction between neurons, and a standard additive white noise.

Based on the Wilson-Cowan formalism, the so-called stochastic rate models [45–47] (see also [48–51]), where the neuron dynamics is described not by the potential of the neuron’s membrane, but by the time-dependent probability of spike generation, can capture the occurrence of the PS regime due to nonlinear phenomenological interaction between populations of neurons. However, the relative freedom of choice of this interaction does not make it possible to obtain either unambiguous and robust biological interpretation or quantitative comparative assessment of results. Nevertheless, stochastic rate models have led to some progress in understanding the oscillatory synchronization of stochastically spiking neurons. For instance, using a very simplified probabilistic neuron model and a functional network connectome extracted from experimental data, the authors of Ref. [51] were able to reproduce a characteristic set of spiking activity patterns obtained by calcium imaging in brain slices with a small number (about 200) of sparsely spiking neurons, i.e. in the case of relatively weak network events similar to population spikes in neuronal networks cultured *in vitro*.

In this paper, we combine with each other the stochastic spontaneous spiking activity of neurons and their deterministic synaptic interaction, which can naturally lead to ‘deterministic’ spikes. We have found that networks of such neurons, both with the metric-free and metric-dependent connectome, demonstrate the regime of repetitive PSs. In contrast to the entirely deterministic model [29, 37], nucleation sites of PSs in the networks with metric-dependent connectome, although do arise, are apparently not stationary. However, due to the difficulty of numerical verification of this assumption, it can only be definitely stated that (i) the number of different nucleation sites at stochastic spontaneous spiking activity of neurons significantly exceeds the corresponding number in the case with a fixed fraction of steady pacemaker neurons [29], and (ii) the relative activity of individual nucleation sites in the stochastic case is on average significantly lower than in the case with steady pacemakers. Finally, the results indicate that the hypothesis of ‘trigger’ neurons, whose activity system-atically precedes the emergence of a PS [52–59], may need further development: even if the probability of spontaneous spike generation was the same for all neurons of the network, repetitive PSs still occurred (cp. [60]). Therefore, the search for such trigger neurons in this case is transferred entirely to the properties of both the concrete implementation of the network connectome and the distribution of amplitudes of synaptic current pulses, excluding intrinsic properties of neurons.

## 2. Neuronal Network Model

The mathematical model of spiking neuronal network consists of three main components: (i) a dynamic model of the neuron that includes a deterministic model of dynamics of the neuron potential, as well as a model of stochastic spontaneous spike generation, (ii) a dynamic model of synaptic interaction between two neurons, and (iii) a static model of the network connectome. In this paper, we consider networks composed only of excitatory neurons, since the presence of inhibitory neurons (with the total fraction of 20% [61, 62]) is not crucial for the considered effects yet masking them and delaying the statistics accumulation [29, 37]. The network model has been studied by numerical simulations performed using custom-made software *NeuroSim-TM* [29] written in the C programming language (the source code and all generated data used for the Figures are available as the Supplementary Material). In particular, ordinary differential equations for the dynamics of neuron potentials and synaptic currents were solved numerically using the standard Euler method with time step Δ*t* = 0.1 ms and with the initial conditions that all neurons had the same ‘resting’ value of the membrane potential, and all synapses had the same initial values of the fractions of synaptic resources. (Note that though setting random initial conditions seems to be a good alternative, it brings additional complexity and hinders the reproducibility of simulations. Therefore, after making sure that the choice between the identical and random initial conditions does not influence essentially the network dynamics, regulating only the moment of the first PS, we have used the simpler variant of identical initial conditions.)

### 2.1. Neuron model

Qualitatively, the neuron model is based on two empirical facts. First, if pulsed stimulation through incoming synapses is cumulatively strong enough for the neuron potential to exceed a threshold value, then the neuron itself generates a spike in response. Second, it has been experimentally established that in a long absence of incoming synaptic current pulses, neurons are able to spontaneously generate spikes on their own [63, 64]. Therefore, the neuron model consists of two complementary parts: 1) a deterministic dynamic model of the neuron potential that allows reproducibly describing spike-based synaptic interaction between neurons, and 2) a stochastic model of spontaneous spike generation. In other words, the neuron model is a leaky input integrator superimposed with a homogeneous Poisson process, modulated by absolute refractory period [65].

Quantitatively, we use the standard Leaky Integrate-and-Fire (LIF) neuron, which has no ability for intrinsic bursting. Subthreshold dynamics of transmembrane potential *V* of such a neuron is described by equation

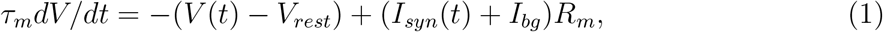

where *V_rest_* is the neuron’s resting potential, *τ_m_* is the characteristic relaxation time of *V* towards *V_rest_*, and *R_m_* is the electrical resistance of the neuron’s membrane. The total incoming synaptic current *I_syn_*(*t*), as a function of time *t*, depends on the choice of the dynamic model of a synapse and the number of incoming synapses. *I_bg_* is an auxiliary constant ‘background’ current, the magnitude of which varies from neuron to neuron. The background currents determine the neuron’s individuality, i.e., the diversity of neuronal excitability, and the fraction of steady pacemakers in the network [63, 64, 66].

When the transmembrane potential reaches a threshold value *V_th_ = V (t_sp_*), it is supposed that the neuron emits a spike, then *V* abruptly drops to a specified value *V_reset_*, *V_rest_* ≤ *V_reset_ < V_th_*, and retains this value during the absolute refractory period *τ_ref_*, then dynamics of the potential is again described by Eq. (1). The result of the LIF neuron dynamics is a sequence of spike generation moments 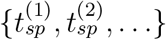.

Numerical values of parameters for the deterministic part of the neuron model [26]: *τ_m_* = 20 ms, *R_m_* = 1 GΩ, *V_rest_* = 0 mV, *V_th_* = 15 mV, *V_reset_* = 13.5 mV, and *τ_ref_* = 3 ms. Finally, Eq. (1) had initial condition *V* (*t* = 0) = *V_rest_* for all neurons.

It is worth explaining the reasons for introducing *V_reset_*, an additional parameter for the LIF neuron. From the computational viewpoint, at *V_reset_ > V_rest_* the neuron’s excitability is increased after the preceding spiking followed by the absolute refractory period. It may facilitate the occurrence of PSs if the incoming synaptic signals are still intense, reducing the time for gathering the PS statistics. From the biological viewpoint, this could also mimic conditions where membrane potentials of neurons fluctuate stationary just a few millivolts below the spiking threshold, as shown by intracellular recordings during the ongoing spiking activity of cortical neurons *in vivo* (sometimes referred as ‘the high-conductance state’ [67]). Importantly, setting *V_reset_ = V_rest_* did not change the simulation results qualitatively.

In this paper, we set *I_bg_* = 0 in Eq. (1) for all neurons in most simulations, except those used in Figs. 1, 7, and 8 for comparison to the main results. A separate consideration of the case *I_bg_* ≠ 0 can be found in Refs. [29, 37], below we have only outlined briefly its key points.

**FIG. 1.**
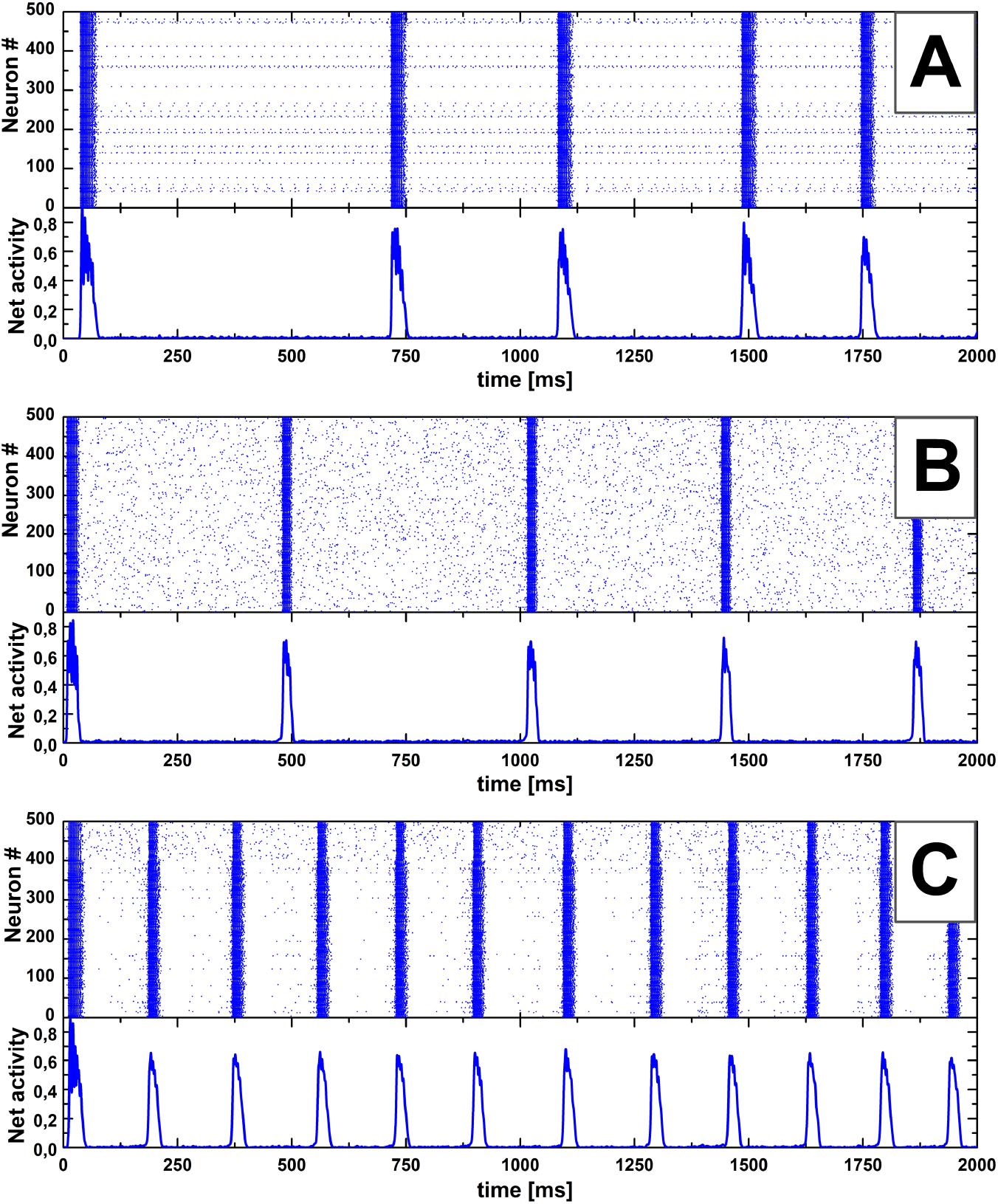
Three examples (A, B, C) of spontaneous population spikes occurring in ‘binomial’ neuronal networks of 500 excitatory LIF neurons with pairwise connection probability *p_con_* = 0.1: raster (top) and network spiking activity, averaged over 2 ms and normalized to the total number of neurons (bottom). Population spikes are the vertical stripes in the raster and the peaks in the activity plot. The networks differ only in neuron parameters (*I_bg_* and/or *p_sp_*), in all the rest (the connectome, distributions of synaptic parameters) the networks have been intentionally made the same. (A) The case with normally distributed values of the background current *I_bg_* and steady pacemakers (3.4% of all neurons, see Sect. 2.1). (B) The case where probability *p_sp_* = 0.0005 of spontaneous generation of a spike per unit time (Δ*t* = 0.1 ms) is the same for all neurons. (C) A hybrid case where 400 neurons have *I_bg_ < I_c_* and *p_sp_* = 0 (i.e. these neurons are definitely not pacemakers) and 100 neurons have *I_bg_* = 0 and *p_sp_* = 0.0005 (i.e. these neurons are stochastic quasi-pacemakers). On the raster graphs, note the pronounced differences in the low-level network activity between population spikes for the considered cases.

**FIG. 2.**
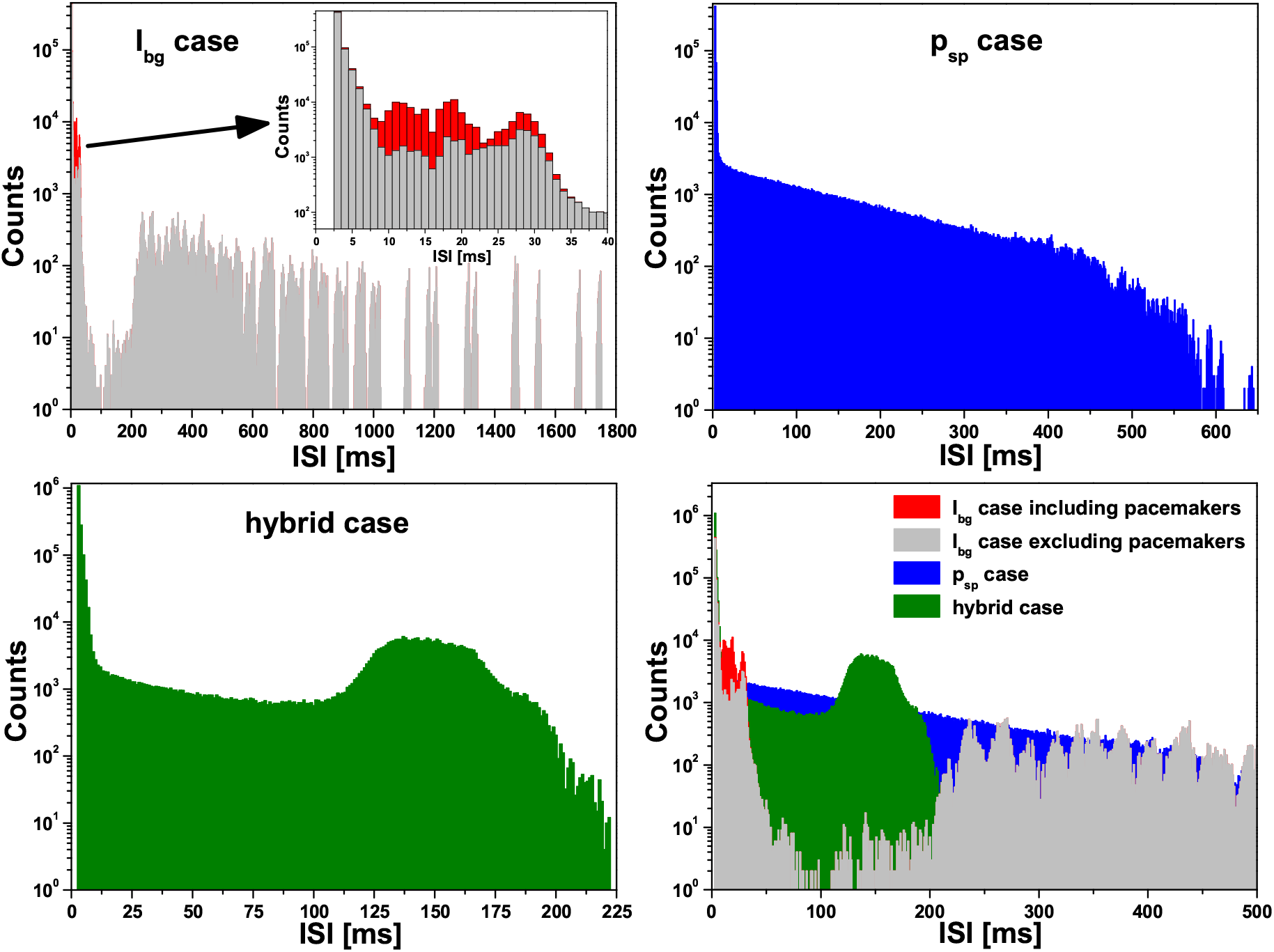
Distributions of Inter-Spike Intervals (ISI) for the ‘binomial’ neuronal networks as those in Fig. 1. Emphasise that these are for ‘elementary’ neuronal spikes, not population spikes. Top left graph: The case of constant background currents resulting in a fixed fraction of pacemaker neurons (’*I_bg_* case’, see also Fig. 1A). The inset shows enlarged view of the region at the origin of the arrow. Gray distribution is for the ISI statistics excluding pacemakers, the red one corresponds to the full ISI statistics. Top right graph: The case of stochastic spontaneous spiking with probability *p_sp_* = 0.0005 (’*p_sp_* case’, see also Fig. 1B). Bottom left graph: The hybrid case, see caption for Fig. 1C. Bottom right graph: Combination of three preceding plots. The ISI distributions clearly show qualitative statistical differences in the three cases.

**FIG. 3.**
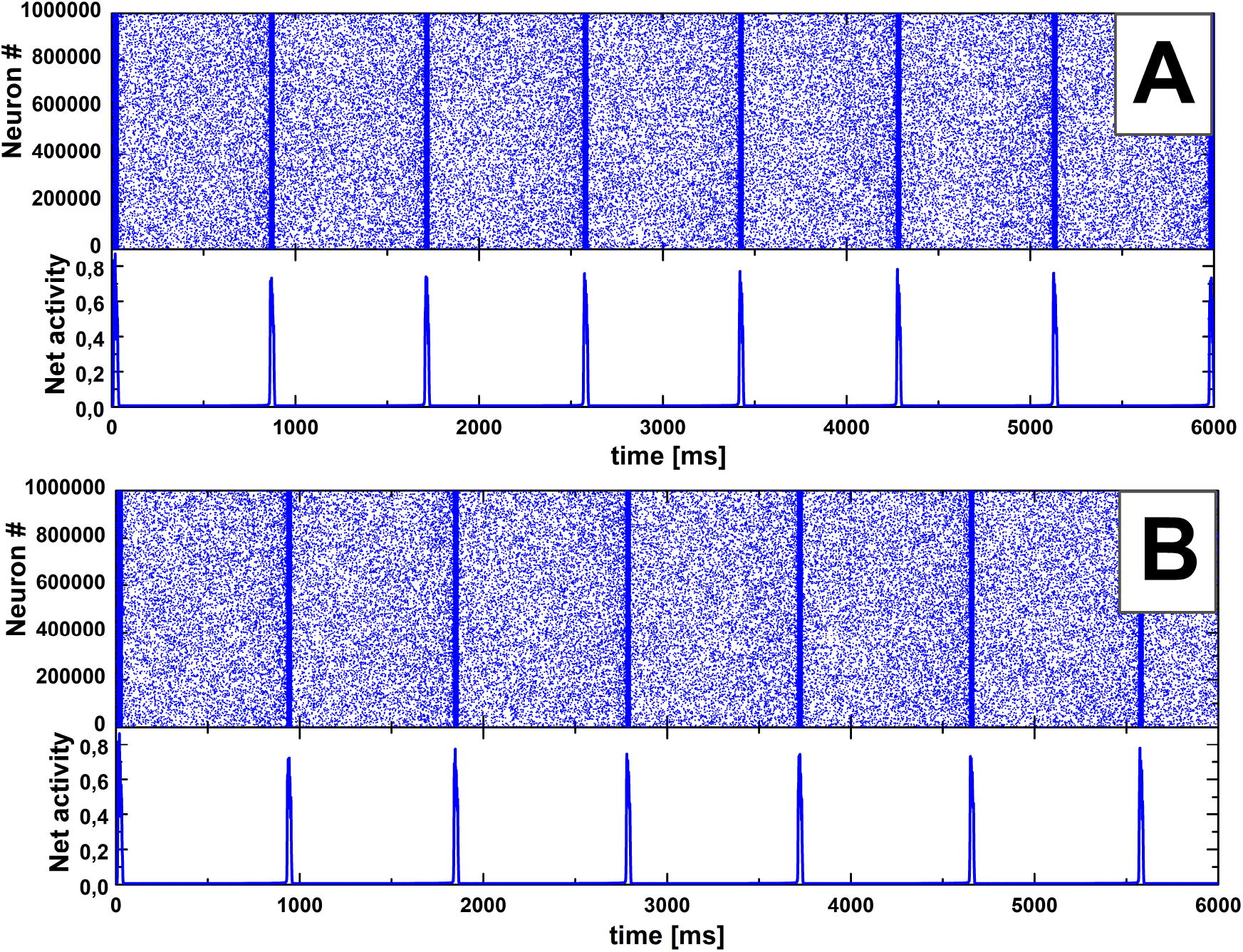
Two examples (A and B) of spontaneous population spikes (PSs) occurring in large ‘binomial’ neuronal networks (with different connectomes etc., unlike the case in Fig. 1) of one million excitatory LIF neurons with *p_con_* = 5 · 10*^−5^* and stochastic spontaneous spiking activity: raster (top) and network spiking activity, averaged over 2 ms and normalized to the total number of neurons (bottom). To reduce the size, the raster graphs show only 0.3% of randomly sieved points (the downsampling algorithm: a random number is generated for each point of the initial raster, and if it is less than the specified value, the raster point is saved on the graph). (A) The case where probability *p_sp_* = 0.0003 of spontaneous spike generation per Δ*t* = 0.1 ms is the same for all neurons. PS periodicity is 855 ± 9 ms (mean ± SD). (B) The case where *p_sp_* value is individual for every neuron, being distributed according to the non-negative and upper-bounded (by 0.001) part of the normal distribution with the mean 0.0003 and standard deviation 0.0001. PS periodicity is 927 ± 12 ms. Note a surprising nearly-periodic regularity (CV ≈ 0.01) of the PSs in both cases.

**FIG. 4.**
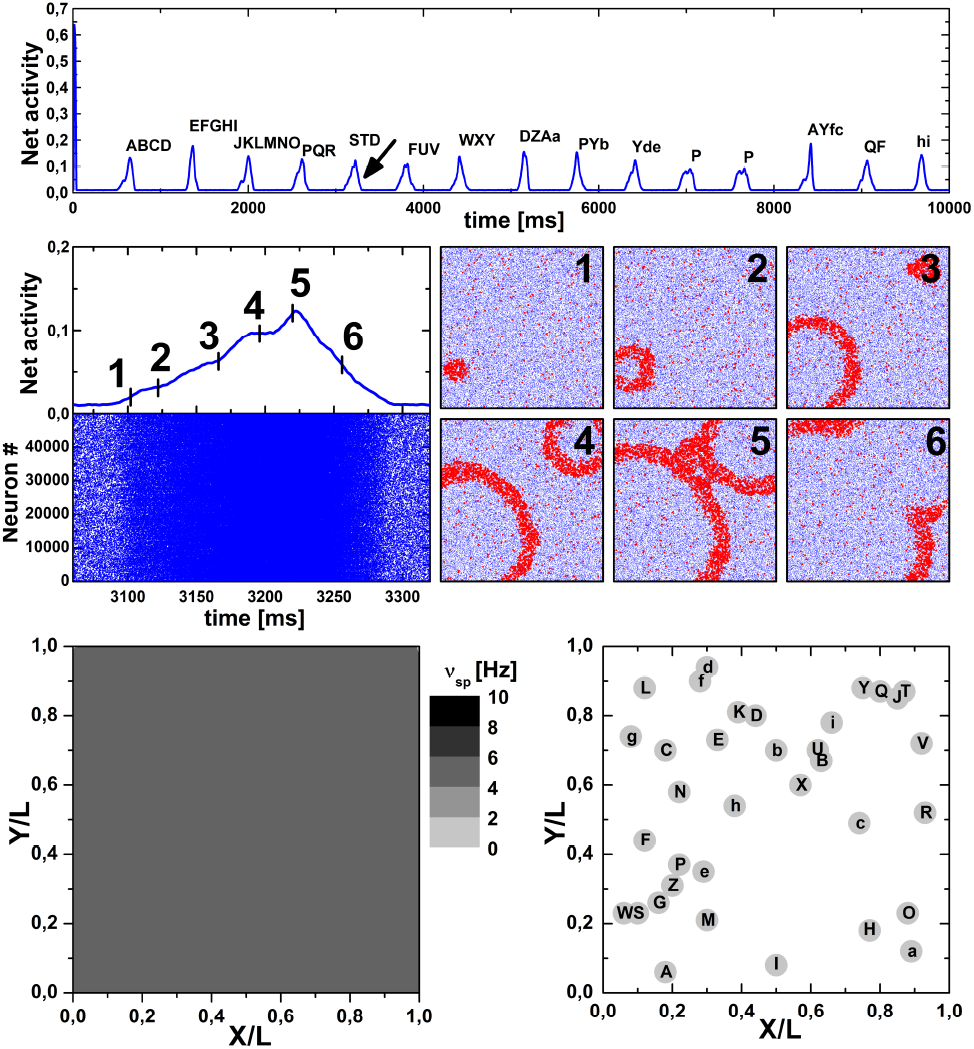
Simulation of spiking activity of a neuronal network consisting of 50 thousand excitatory LIF neurons statistically uniformly distributed over the square *L* × *L*. Synaptic connections have been formed with probability *p_con_* = exp(−*r/λ*) + *p*_min_*θ*(*r* − *r*_0_), see Eq. (15), where *λ* = 0.01*L*, *p*_min_ ≈ 3 · 10*^−5^*, and *r*_0_ ≈ 0.1*L*. This gives 32 ± 6 (mean ± SD) outgoing connections per neuron. All neurons have the same value *p_sp_* = 0.0005 of the probability of spontaneous generation of a spike per unit time (Δ*t* = 0.1 ms). Upper graph: Network spiking activity, averaged over 2 ms and normalized to the total number of neurons, during 10 seconds of the simulation. Each population spike is denoted by a sequence of Latin letters (uppercase and lowercase letters are not the same), indicating the activation sequence of nucleation sites underlying that population spike. Middle graph: LEFT: Network activity (top) and its raster (bottom) during the population spike marked by the arrow in the upper graph. RIGHT: Six snapshots of the instantaneous spatial spiking activity of neurons for the corresponding moments (labeled by the numbers from 1 to 6) of the population spike. Each frame corresponds to the whole area *L* × *L*. Blue dots depict inactive neurons and red dots highlight active neurons. Bottom graph: LEFT: Spatial distribution of neurons with a certain value of spontaneous spiking rate *ν_sp_* ≈ *p_sp_/*Δ*t*. As for this simulation all neurons have the same *p_sp_*, the distribution is absolutely uniform. RIGHT: Schematic reconstruction of the spatial pattern of emergent nucleation sites (depicted by filled gray circles) for all population spikes shown in the upper graph. The graphs show 15 PSs and 35 nucleation sites with 11 primary ones (labeled as A, E, J, P, S, F, W, D, Y, Q, h), with only 2 of them (A, P) activated more than once.

**FIG. 5.**
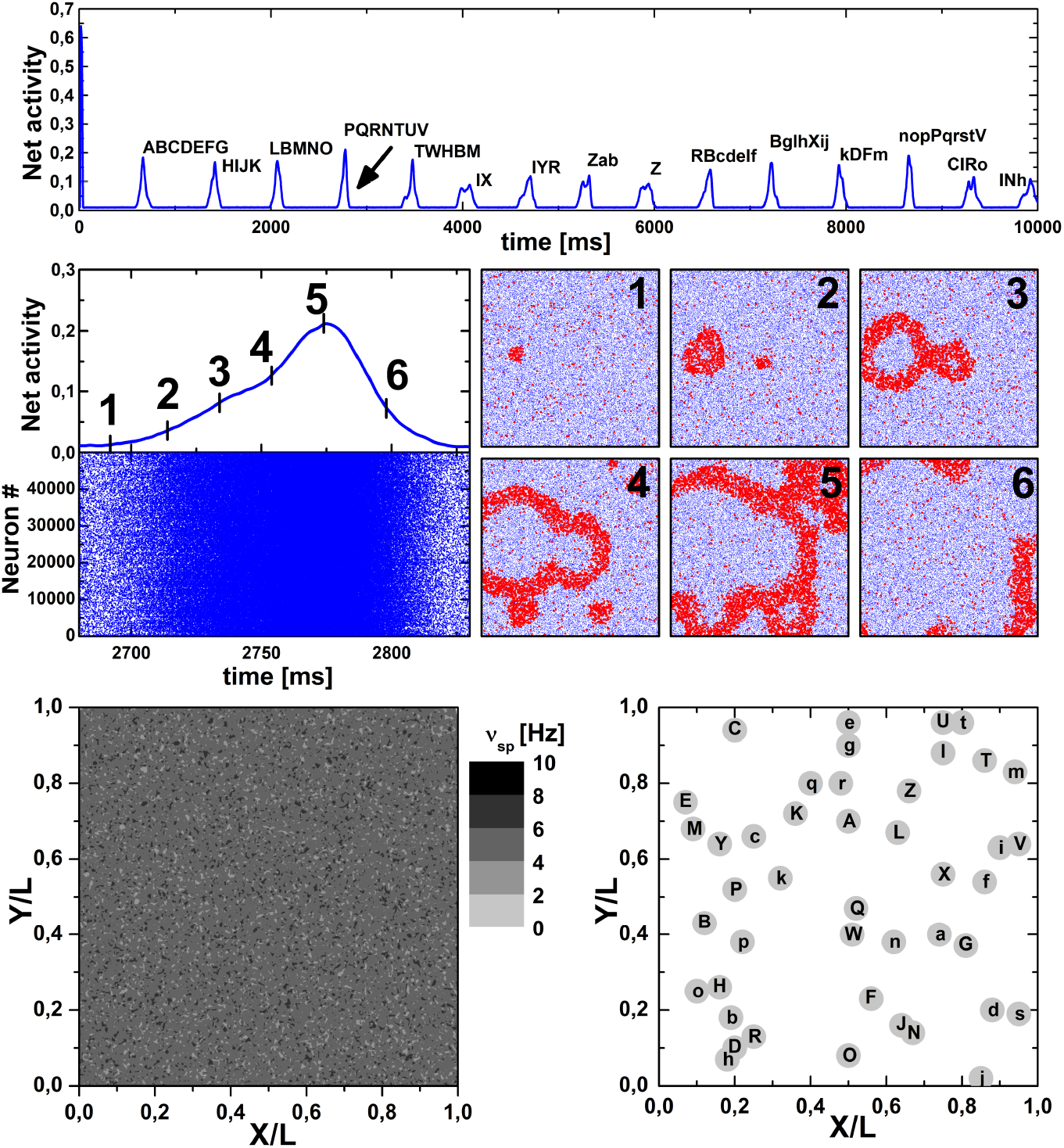
Simulation of spiking activity for the same neuronal network as in Fig. 4 except the set of *p_sp_* values for the neurons: unlike the former case in Fig. 4, where *p_sp_* is the same for all neurons, now *p_sp_* values are distributed among neurons according to the non-negative and upper-bounded (by 0.001) part of the normal distribution with the mean 0.0005 and standard deviation 0.0001. All graphs have the same meaning as those in Fig. 4. Note a dispersed spatial distribution of neurons with a certain *ν_sp_* value and a different spatial pattern of emergent nucleation sites (left and right bottom graphs, respectively) compared to the case in Fig. 4. One can see 15 PSs and 44 nucleation sites with 12 primary ones (A, H, L, P, T, I, Z, R, B, k, n, C), with only 2 of them (I, Z) activated more than once.

**FIG. 6.**
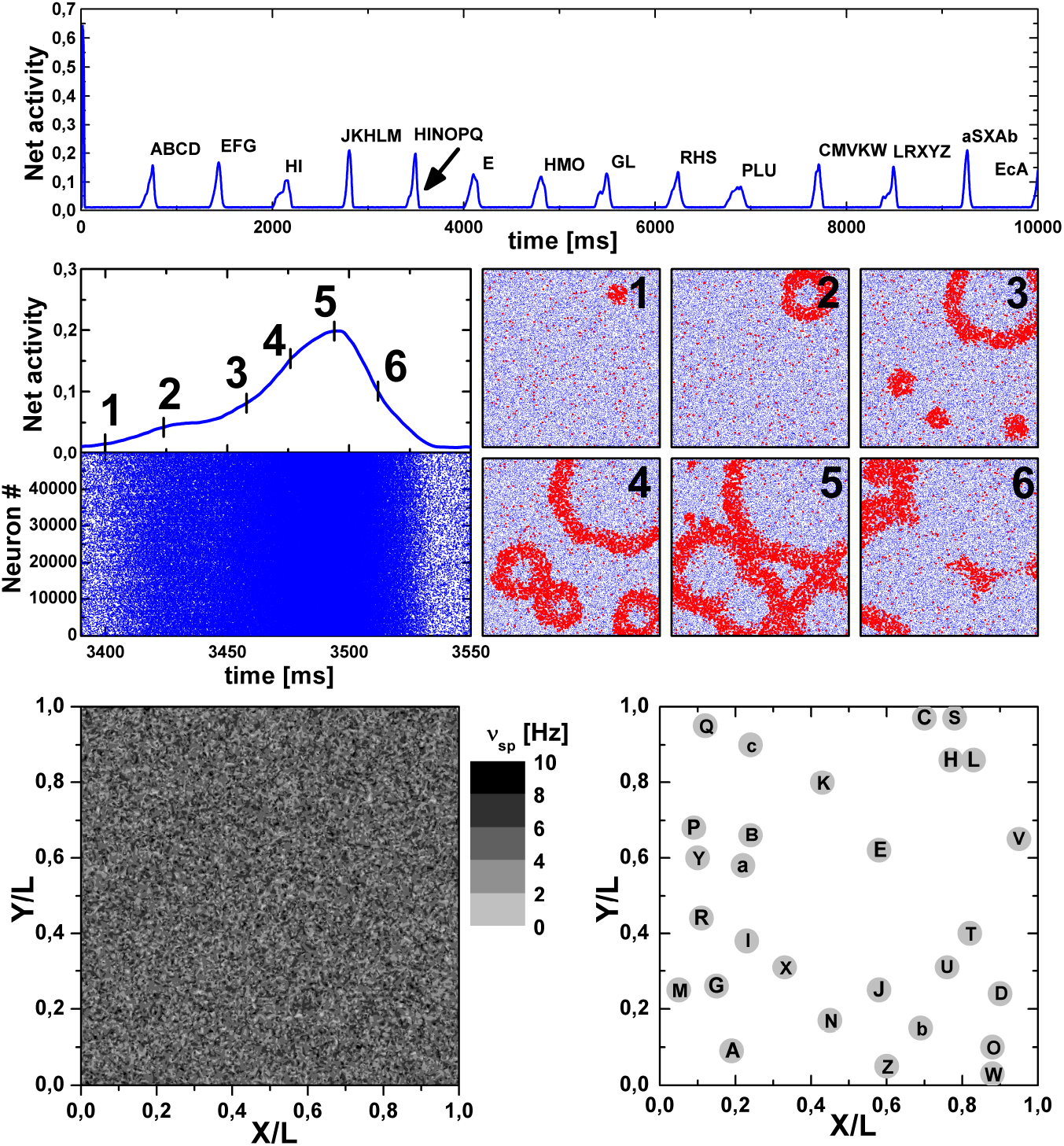
Simulation of spiking activity for the same neuronal network as in Fig. 5 except a different set of *p_sp_* values for the neurons: now *p_sp_* values are distributed with the standard deviation 0.0002, all the rest is the same as in Fig. 5 (each graph also has the same meaning as before). Note the increased dispersion of the spatial distribution of neurons with a certain *ν_sp_* value and a new spatial pattern of emergent nucleation sites (left and right bottom graphs, respectively) compared to the cases in Figs. 4 and 5. In this case, there are 14 PSs and 28 nucleation sites with 10 primary ones (A, E, H, J, G, R, P, C, L, a), again, with only 2 of them (E, H) activated more than once.

**FIG. 7.**
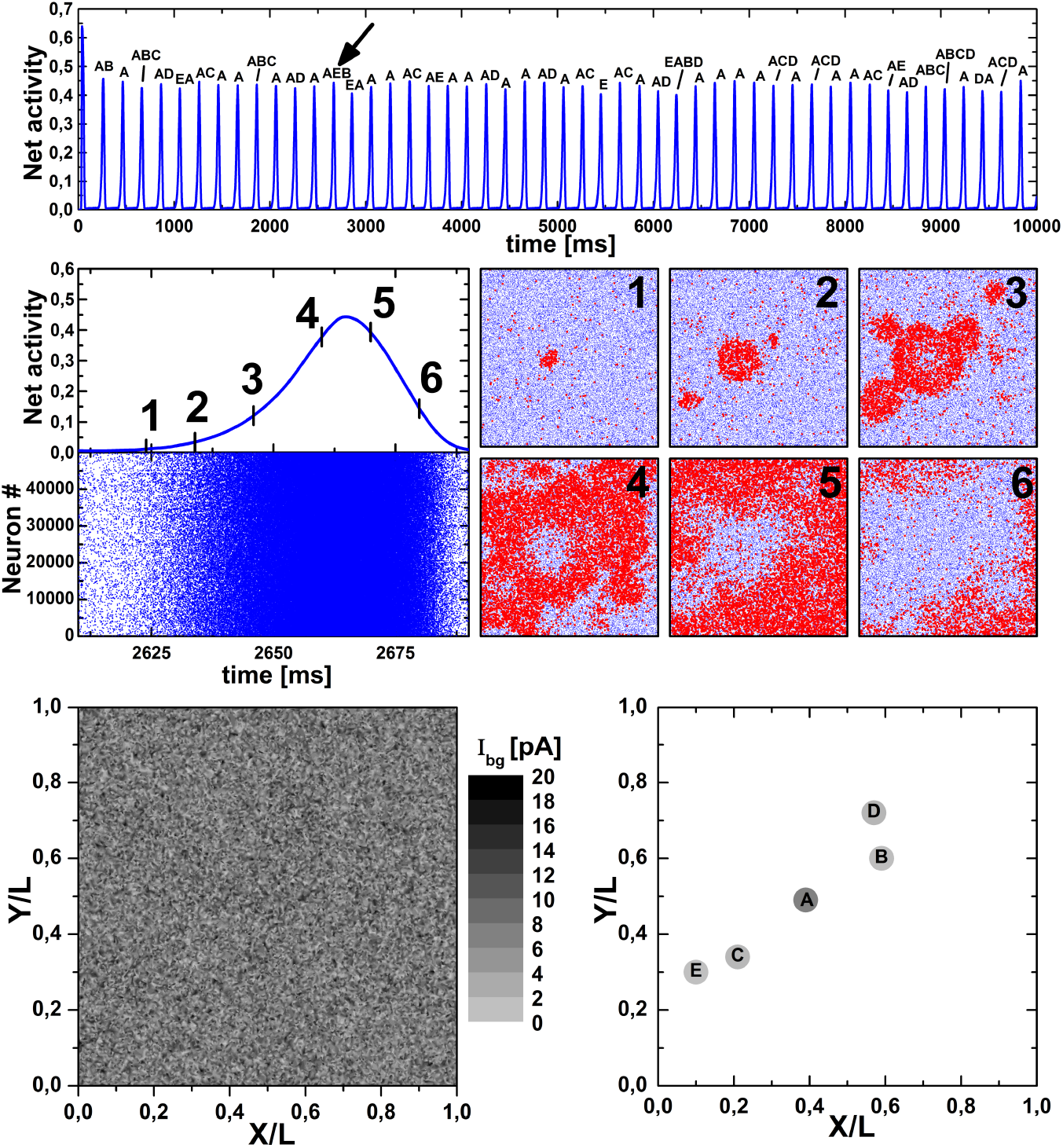
Simulation of spiking activity for the same metric neuronal network as in Fig. 4 except that instead of stochastic spontaneous spiking with probability *p_sp_* all neurons have normally distributed values of the background current *I_bg_* (as in Fig. 1A), resulting in some fraction of steady pacemakers (3.4% of all neurons, see Sect. 2.1). All graphs have the same meaning as those in Fig. 4, except for the bottom left graph, which now shows spatial distribution of neurons with a certain value of *I_bg_*. The pacemaker neurons have *I_bg_ > I_c_* = 15 pA. In a drastic contrast to the case of stochastic spontaneous spiking (Figs. 4–6), here 49 population spikes shown on the top graph occur from only five steady nucleation sites depicted on the bottom right graph by filled gray circles, with the color depth reflecting their relative activation rate.

**FIG. 8.**
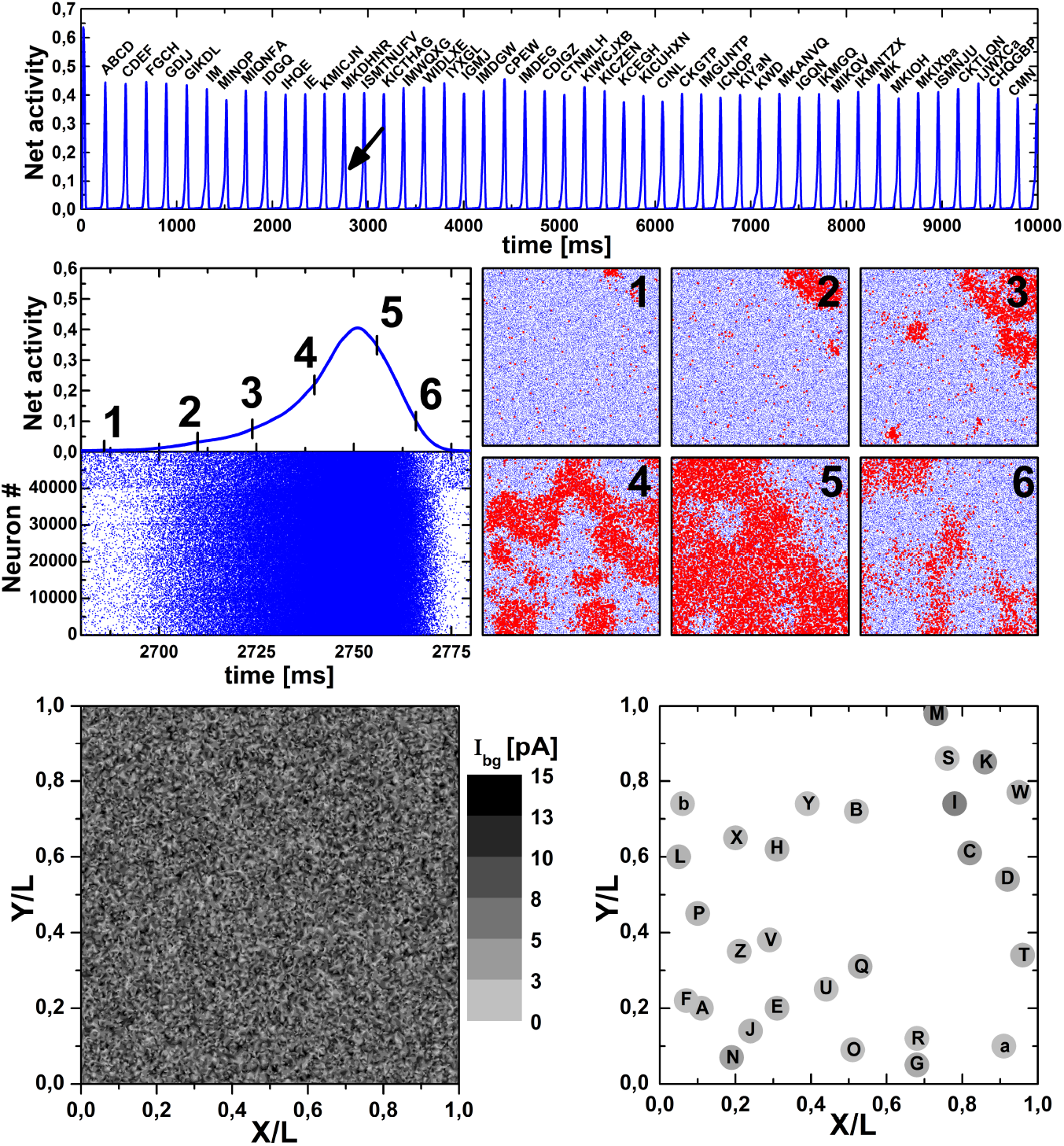
The hybrid case for metric networks: simulation of spiking activity for the same neuronal network as in Fig. 4 except that 80% of neurons have *I_bg_ < I_c_* and *p_sp_* = 0 (i.e. these neurons are not pacemakers) and 20% of neurons have *I_bg_* = 0 and *p_sp_* = 0.0005 (i.e. these neurons are stochastic quasi-pacemakers), just like in Fig. 1C for the binomial network. All graphs have the same meaning as those in Fig. 7. Note that despite a large number of nucleation sites (similarly to the cases in Figs. 4–6) a substantial fraction of them is relatively stable, i.e. population spikes (PSs) occur more than once from some nucleation sites during the simulation. In particular, 47 population spikes shown on the top graph occur from only eight ‘primary’ nucleation sites (A, C, F, G, I, M, K, W) of 28 sites in total, with five sites (C, G, I, M, K) activated more than once.

If a neuron has the value of *I_bg_* that exceeds a critical value *I_c_* = (*V_th_* − *V_rest_*)*/R_m_*, then this neuron is a steady pacemaker (pm), i.e. it is able to emit spikes periodically with frequency

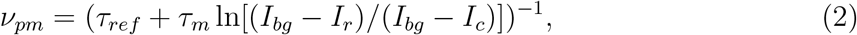

where *I_r_* = (*V_reset_* − *V_rest_*)*/R_m_*, in the absence of incoming signals from other neurons. The above specified parameters of the neuron model give the critical current value *I_c_* = 15 pA and *I_r_* = 13.5 pA.

In turn, *I_bg_ < I_c_* leads to an increase of depolarization of the neuron’s potential to some asymptotic subthreshold value, i.e. to the effective renormalization of the neuronal resting potential, 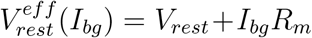, if the neuron is uninfluenced through incoming synapses for a relatively long time.

Provided that the background current values are distributed among neurons according to the non-negative and upper-bounded part of the normal distribution, with the mean *μ* and standard deviation *σ*, the relative fraction of steady pacemakers in an ensemble (in fact, in a network) of *N* neurons is explicitly given by formula [37] (see also [68])

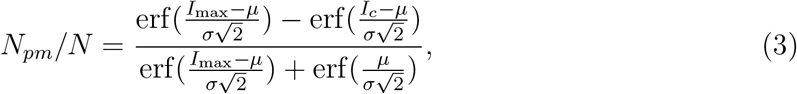

where *I*_max_ is the upper value of the background current. In all simulations the inequalities *σ < μ < I_c_* are assumed, resulting in *N_pm_* ≪ *N*. For instance, with *I*_max_ = 20 pA, *μ* = 7.7 pA and *σ* = 4.0 pA used in most simulations, one gets the fraction of pacemakers *N_pm_/N* = 3.4% with the maximal *ν_pm_* value 121 Hz.

To initiate spiking activity of the network neurons in the case of the absence of background currents (*I_bg_* ≡ 0), we set a probability *p_sp_* of spontaneous generation of a spike per unit time (i.e., within elementary time step Δ*t* = 0.1 ms): for each neuron at every time step, a random real number from zero to one is generated and compared with *p_sp_*. If this number is less than *p_sp_*, the potential of the neuron is set equal to the threshold value, i.e. the neuron emits a spike (provided that this neuron has not emitted spike a little earlier and is not in the refractory state). The presence of the absolute refractory period *τ_ref_* decreases the probability of spontaneous spike generation per Δ*t* to the average value [65]

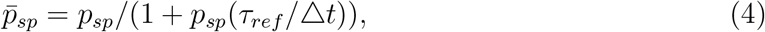

which determines the average spiking rate 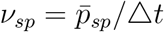.

The normalized spiking activity of the neuronal network is defined as a ratio of the number of neurons that emitted spikes during some averaging interval *t_avg_* to the total number *N* of network neurons:

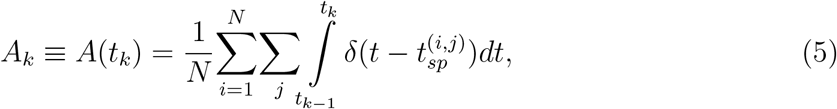

where *t_k_ = kt_avg_*, *k* = 1, 2,… is the sequence of natural numbers, *t*_0_ = 0, and 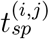 is the moment when *i*-th neuron generates its *j*-th spike. In the limiting case *t_avg_* = Δ*t* for *N* disconnected, independently spiking neurons with the same initial condition, one can obtain an exact linear recurrent sequence for *A_k_* analytically (with *n_ref_ = τ_ref_ /*Δ*t*) [65]:

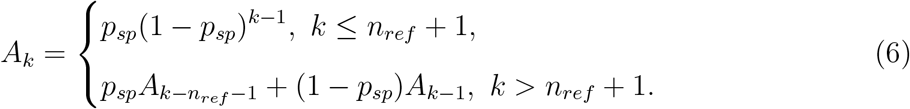

Here *A_k_* has an accurate meaning of spike generation probability for a statistical ensemble of *N* neurons at *k*-th time step. At *k* → ∞ this converges to *A_∞_ = p_sp_/*(1 + *p_sp_n_ref_*), i.e. to the average value 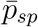 determined by Eq. (4). Note that Eq. (6) describes transient damped oscillations in the general case [65].

While simulating relatively large-scale networks with more than a few hundred neurons, it is practical to set *t_avg_* > Δ*t*, typically *t_avg_* ∼ *τ_ref_*. In the case of non-predefined *t_avg_* value, the normalized average activity of *N* disconnected stochastic neurons is *ν_sp_t_avg_*. Based on extensive experimental data on spontaneous neuronal activity both *in vitro* and *in vivo* [69–78], we took *p_sp_* = 0.0005 (or *ν_sp_* ≈ 5 Hz) so that the baseline level of asynchronous spontaneous network activity at *t_avg_* = 2 ms was equal to a few percent of the total number of neurons, *ν_sp_t_avg_* ≈ 0.01.

### 2.2. Synapse model

The dynamic model of the synapse incorporates a phenomenological model of short-term synaptic plasticity [25, 26, 79, 80] (cp. [81, 82], reviewed in [83, 84]), which is consistent with independent experiments [85, 86] (see also [87, 88]). For the sake of integrity, along with the description for excitatory synapses, we also concisely provide the specifics of the inhibitory synapses within the model.

A single contribution to the incoming synaptic current is determined as

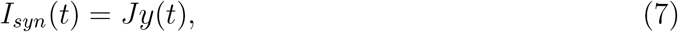

where *J* is the maximal amplitude of synaptic current, the sign and magnitude of which depend on the type of pre- and postsynaptic neurons (i.e., whether the neuron is excitatory or inhibitory), and *y*(*t*) is a dimensionless parameter, 0 ≤ *y* ≤ 1, the dynamics of which is determined by the following system of equations [26]:

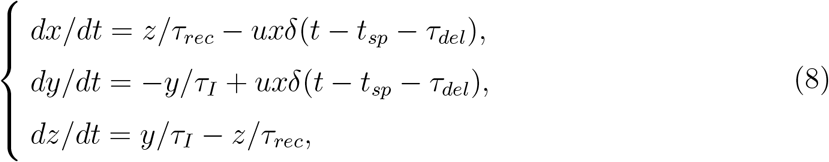

where *x*, *y* and *z* are the fractions of synaptic resources in the recovered, active and inactive states, respectively, *x + y + z* = 1, *τ_rec_*, *τ_I_* are the characteristic relaxation times (*τ_rec_* ≫ *τ_I_*), *δ*(…) is the Dirac delta function, *t_sp_* is the moment of spike generation at the presynaptic neuron, *τ_del_* is the spike propagation delay (*τ_del_* ≡ 0 for metric-free networks; for metric networks, see Eq. (14) below), and *u* is the fraction of recovered synaptic resource used to transmit the signal across the synapse, 0 ≤ *u* ≤ 1. For the outgoing synapses of inhibitory neurons, the dynamics of *u* is described by equation [25, 26]

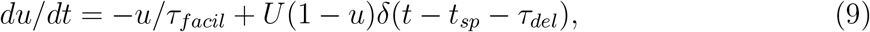

where *τ_facil_* is the characteristic relaxation time, and 0 < *U* ≤ 1 is a constant parameter. For the outgoing synapses of excitatory neurons, *u* remains constant and equals to *U*.

Qualitatively, the dynamics following from Eqs. (8) implies a reversible depression of synaptic transmission due to the depletion of synaptic resources at intense incoming spike train, while the additional Eq. (9) for inhibitory synapses enables them to overcome the depression and to facilitate the transmission [80, 89, 90].

In numerical simulations all synaptic parameters, except *τ_I_*, were taken from the fixed-sign parts of the normal distributions with the mean values described below, i.e. each synapse had its own unique values of these parameters. The standard deviations for all distributed parameters equal half of their mean values; see more detail in Refs. [29, 37].

Numerical values of parameters for the synapse model [26] (see also [80]): *τ_I_* = 3 ms, mean values for the normal distributions were *τ_rec,ee_ = τ_rec,ei_* = 800 ms, *τ_rec,ie_ = τ_rec,ii_* = 100 ms, *τ_facil,ie_ = τ_facil,ii_* = 1000 ms, *J_ee_* = 38 pA, *J_ei_* = 54 pA, *J_ie_ = J_ii_* = −72 pA, *U_ee_ = U_ei_* = 0.5, *U_ie_ = U_ii_* = 0.04. Here, the first lowercase index denotes the type (*e* = excitatory, *i* = inhibitory) of the presynaptic neuron, and the second index stands for the type of the postsynaptic neuron. Initial conditions for Eqs. (8) were the same for all synapses: *x*(*t* = 0) = 0.98, *y*(*t* = 0) = 0.01, *z*(*t* = 0) = 0.01.

### 2.3. Network connectome model

Despite that the main focus of this study is on the networks with spatially-dependent connectome (‘metric networks’) of small-world type, we preliminary consider a simpler case of abstract ‘metric-free’ networks with the binomial distribution of connections between neurons. (Hereafter, the distribution of connections means the degree distribution in terms of network science: the degree of a neuron is the number of connections it has to other neurons. The connection strength is not implied anywhere in the description of the network connectome model.) Even relatively small metric-free networks can exhibit irregular spontaneous PSs [26], making this network type quite helpful for studies of statistical properties of the PS regime.

Specifically, for a metric-free ‘binomial’ network, its connectome is a directed graph, the nodes of which are point neurons, with the binomial distribution of connections between nodes. When creating a connectome realization, for each directed pair of neurons, a random real number *ξ* is generated in the range from zero to one and compared with a given spatially-independent probability *p_con_* of the formation of unilateral synaptic connection between two neurons. If *ξ* ≤ *p_con_*, the connection is formed; otherwise, it is not formed. To reduce the complexity of the resulting model, in what follows, we assume that the formation of autaptic connections (i.e., self-connections, see [91]) is not allowed. Then, for a network of *N* neurons, one gets the binomial distribution for the number *n* of outgoing (or, equally, incoming) connections per neuron, 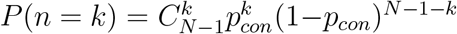, with the mean value 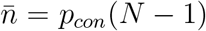 and variance 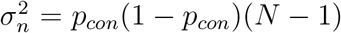. The average number of network connections is 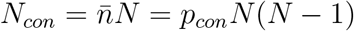. Since usually *N* >> 1, in what follows, we will always count (*N* −1) ≈ *N*. Also note that the coefficient of variation 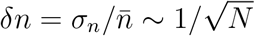, so for *N* >> 1 deviations from the mean value are insignificant. In most numerical simulations we studied networks of *N* = 500 excitatory neurons with *p_con_* = 0.1 and 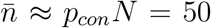 (see Fig. 1). In addition, we performed two large-scale simulations of binomial networks of *N* = 10^6^ excitatory neurons (Fig. 2) with *p_con_* = 5 · 10*^−5^* resulting in the same 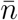 value. The connectome of each network was created (or restored from a file) before starting the dynamic simulation and remained unchanged, i.e. static, during the simulation.

In the case of spatially-dependent network topology, in order to set the connections between neurons, one needs to assume (i) how neurons are distributed in space, i.e., to set their spatial coordinates and a metric distance between every two neurons, and (ii) how the probability of forming a unilateral connection depends on the distance between neurons.

In accordance with the technique for preparing two-dimensional neuronal cultures [7], we assume that *N* point neurons are *uniformly* distributed over the square area *L* × *L*. Then the probability density *P* (*r*) of detecting two neurons at distance *r* from each other is given by (*r* is expressed in units of *L*) [29]

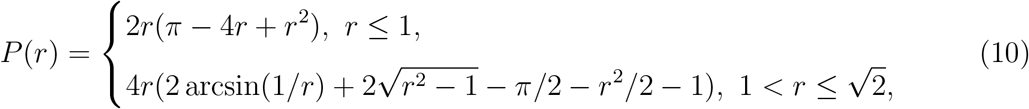

such that 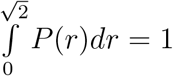. Function *P* (*r*) has a single maximum at 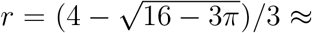 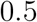.

Next, we assume that the probability of forming a unilateral connection between every two neurons *decreases exponentially* with increasing distance *r* between them [92–97] (cp. [27, 98]),

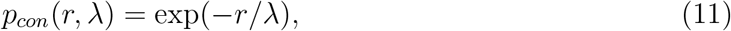

where *λ* is a characteristic connection length, which is also expressed in units of *L*. The formula (11) is slightly idealized; its actual modification follows below (see Eq. (15)).

With a given function *p_con_*(*r, λ*), the average number of connections in a network of *N* neurons is 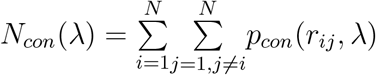, where *r_ij_* is the distance between the *i*-th and *j*-th neurons. If neurons are uniformly distributed over the square area, *r_ij_* is a random value with the distribution density *P* (*r*) given by Eq. (10).

In addition, as the square area is a convex set of points, we have assumed that the connections between neurons can be modeled by straight line segments. Importantly, the connections do not cross boundaries of the square. Due to this, neurons in the vicinity of the boundaries have fewer connections.

With the above assumptions, the resulting distribution of connection lengths is given by the product *p_con_*(*r, λ*)*P* (*r*), which reaches its maximum at *r* ≈ *λ* for *λ* ≲ 0.1 (see Supplementary Material in [29] for more details).

The average number of network connections can be expressed as 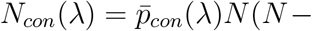 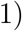, where the space-averaged probability

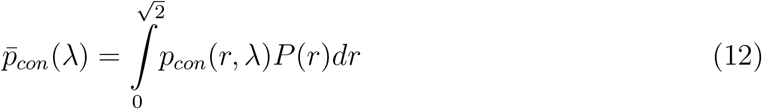

increases monotonically with *λ*, asymptotically reaching unity at infinity. For *p_con_*(*r, λ*) determined by Eq. (11), at *λ* ≪ 1 one gets 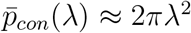. (For the whole range of *λ*, the approximate analytical expression for 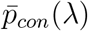 is given in [29].) In turn, the average number of outgoing (or incoming) connections per neuron is 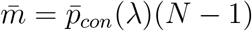.

In addition, it is useful to determine explicitly the average fraction of network connections with the lengths longer than or equal to *l*, where 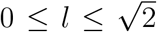. Denoting that fraction as *n_con_*(*l*), one gets

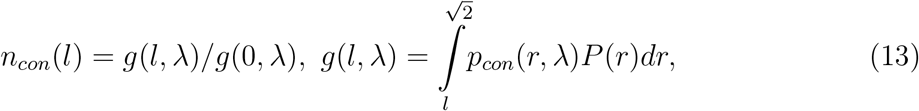

with 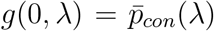. The dependence *n_con_*(*l*) is shown on the right graph in Fig. S1 in folder ‘Additional Simulations’ of the Supplementary Material.

Finally, due to the fact that every connection between neurons has a metric length, there exist spike propagation delays. Assuming a constant speed of spike propagation along connections (essentially, axons), the delays are calculated by formula

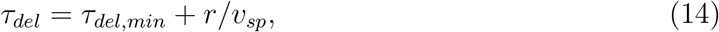

where *τ_del_* is the total spike-propagation delay for a connection of length *r*, *τ_del,min_* is the minimal delay same for all neurons, and *v_sp_* is the constant speed of spike propagation. Note that the distribution of ‘axonal’ delays (14) is also determined by the product *p_con_*(*r, λ*)*P* (*r*). To implement the network connectome models above, a random number generator (RNG) is necessarily required. We show below that in the spatially-dependent case RNG may lead to a systematic bias for the fraction of long-range connections. As mentioned earlier, the described model was implemented in the C programming language, which had been chosen because of comparatively fast execution of a program code. The standard RNG in C can generate a random integer number within the range from 0 to some predetermined value *RAND MAX*, which may depend on the operating system but cannot be smaller than 32767. That integer number can be then transformed into the real one by normalizing it by *RAND MAX*, resulting in the pseudo-uniform distribution from 0 to 1 that is used to generate neuronal connections by comparing the random numbers with the probability (11). However, as the smallest nonzero random number is *p*_min_ = 1*/RAND MAX*, there is a systematic bias for nonzero probability values smaller than this limit: these values are compared with zero and always lead to the connection formation. As a result, the actually implemented model is as follows:

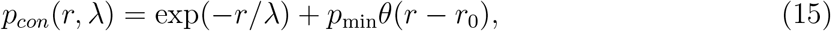

where *r*_0_ = *λ* ln(1*/p*_min_) = *λ* ln(*RAND MAX*), and *θ*(*…*) is the unit step function: *θ*(*x*) = 1 for *x* > 0 and *θ*(*x*) = 0 for *x* ≤ 0. If *RAND MAX* turns to infinity, the second term in (15) vanishes and we return to the initial model (11). Given *RAND MAX* = 32767 and *λ* = 0.01, one gets *p*_min_ ≈ 3 · 10*^−5^* and *r*_0_ ≈ 0.1. Note that for the large-scale binomial networks with *p_con_* = 5 · 10*^−5^*, no statistical deviation in the number of network connections was detected.

Numerical values of parameters for the connectome of metric networks were as follows [29, 37]: *N* = 50000, *λ* = 0.01*L*, *τ_del,min_* = 0.2 ms, and *v_sp_* = 0.2 *L*/ms with *L* = 1 mm by default.

In all spatial simulations presented in the Results section, we used the connectome model with *p_con_*(*r, λ*) given by Eq. (15). For comparison, we have performed some key simulations with network connectomes generated separately using advanced RNGs in MATLAB and in the NumPy library for the Python programming language. These connectomes fully correspond to the initial model (11) and, essentially, are based on the same set of spatial coordinates of neurons as for the original connectomes with (15).

In particular, the average number of network connections for (15) is 4.8% greater than that for (11), with most of these ‘additional’ connections being long-range ones. In turn, the average number of outgoing connections per neuron is 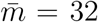 for (15) vs 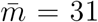 for (11), with the unchanged standard deviation (= 6). Thus, effectively, every neuron acquires one additional long-range connection.

As the fraction of additional connections is quite small, the simulation results have been qualitatively unchanged (see Figures S1–S5 in folder ‘Additional Simulations’ of the Supplementary Material). The only noticeable stable effect was that the absence of additional long-distance connections increases (in about two times) the duration of population spikes and decreases (also in about two times) their amplitudes in the case where spontaneous spiking of neurons occurs deterministically due to steady pacemakers [29, 37], i.e., neurons with supercritical *I_bg_* values (see Eq. (1)). In the case of stochastic spontaneous spiking, the impact of long-distance connections on the formation of unstable nucleation patterns seems to be generally less pronounced due to the lower excitability of neurons in response to a given stimulation, as in this case *I_bg_* = 0 for every neuron.

Finally, it is worth noting that the considered metric networks with *N* = 50000 and *λ* = 0.01*L*, regardless of the particular definition ((11) or (15)) for *p_con_*(*r, λ*), belong to the small-world network type [99–102], with the following mean values of the clustering coefficient (CC) and the shortest path length (SPL): CC ≈ 0.13 and SPL ≈ 4 for the model (15), CC ≈ 0.15 and SPL ≈ 11 for the initial model (11), and 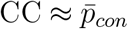 and SPL ≈ 3 for the metric-free binomial networks of the same size and with the same mean degree (for details, see subfolder ‘CC and SPL computation’ in ‘Additional Simulations’ of the Supplementary Material). Based on these numbers, the ‘small-world-ness’ introduced in [100] drops more than twice (from about 103 to 45) with replacing (15) by (11). It is worth noting that the above values of CC and SPL for the metric networks generated using Eq. (15) (but not (11)) are consistent with the independent estimates (see Fig. 3a in [103]).

## 3. Results

Provided stochastic spontaneous activity of neurons, the regime of repetitive population spikes (PSs) has been found both in the metric-free ‘binomial’ networks (Figs. 1 and 3) and in the metric ones (Figs. 4–6).

In contrast to the binomial networks driven by constant background currents, where during the intervals between PSs the spiking activity is mainly determined by steady pacemakers and is therefore a sum of nearly periodic signals, in the network of neurons with stochastic spontaneous activity the intervals between PSs are filled by predominantly stochastic activity (see the graphs A and B in Fig. 1). In addition, we have considered the case of a ‘hybrid’ network, where most of the neurons have subcritical background currents (i.e., these neurons are not pacemakers) and a smaller fraction has no background currents, but instead these neurons can spontaneously emit spikes with the equal probability *p_sp_* per time step. As it turned out, PSs also occurred in this case (Fig. 1 C), and even more intensely than in purely ‘deterministic’ (Fig. 1 A) and purely ‘stochastic’ (Fig. 1 B) cases. Specifically, for the binomial networks demonstrated in Fig. 1 the PS periodicity was as follows: 555 ± 356 ms (mean ± SD) with coefficient of variation CV = 0.64 in the *I_bg_* case (Fig. 1 A), 473 ± 63 ms with CV = 0.13 in the *p_sp_* case (Fig. 1 B), and 169 ± 17 ms with CV = 0.10 in the hybrid case (Fig. 1 C).

Note that the very possibility of an occurrence of recurring PSs is embedded in the model of short-term synaptic plasticity, provided that the stationary network activity between PSs is low enough to allow the synaptic resources to be replenished. This condition is well fulfilled in the case of constant background currents, where the majority of neurons are silent most of the time and PSs are triggered due to stimulating activity of steady pacemakers when the efficiency of synaptic connections between most neurons is sufficiently high (see Fig. 1 in [26] and Fig. 2 in [29]). In the case of the same probability *p_sp_* of spontaneous spiking, when all neurons are independently active and equally excitable, the required balance of available synaptic resources and, as a consequence, the emergence of repeating PSs are not guaranteed. In turn, the hybrid case was designed to explore the influence on PSs by the primal spiking activity that was changed from deterministic and relatively high-rate activity of a small set of steady pacemakers to completely stochastic and low-rate activity of a substantially (in several times) larger set of quasi-pacemakers.

Even more striking differences between these three cases (i.e. *I_bg_*, *p_sp_* and the hybrid ones) are revealed by the distribution of intervals between successive spikes (or action potentials so as not to confuse with PSs) for every neuron of the network. The corresponding distributions of inter-spike intervals (ISI) are shown in Fig. 2. In fully deterministic *I_bg_* case the ISI distribution is comparatively broad and ragged. It exhibits multiple peaks, which are conserved even after excluding the activity of pacemaker neurons from the ISI sampling. The peaks at the beginning (shown on the separate inset in Fig. 2), in the range up to 35 ms, are more regular than subsequent ones and are most sensitive to pacemaker activity. Notably, these peaks could be attributed to some form of stochastic resonance (see Discussion below). In turn, fully stochastic *p_sp_* case exhibits almost monotonically descending ISI distribution, with small peaks only at the large ISI values. Finally, in the hybrid case the ISI distribution starts as in the *p_sp_* case but further it has a large peak within 135-165 ms that is about the mean PS periodicity. As shown further, the qualitatively different ISI distributions for the binomial networks partially conserve for the spatial networks.

Importantly, the PS regime in the binomial networks is easily scalable with the number of network neurons by means of keeping the same mean number of connections per neuron. Indeed, for relatively large-scale binomial networks of one million excitatory LIF neurons the PSs occurrence is conserved and, moreover, it becomes even more regular, nearly periodic (especially in the case where *p_sp_* value is the same for all neurons), see Fig. 3. As shown further, two-dimensional networks having a different type of network topology can also exhibit similar temporal regularity of PSs coexisting with highly irregular spatial dynamics (cp. [15]).

For the metric networks with stochastic spontaneous activity of neurons, compared to the case with a fraction of steady pacemakers, PSs also arise from nucleation sites (Figs. 4–6). However, the number of nucleation sites is significantly (in many times) larger, and their relative activation rates are on average significantly smaller, than in the case of steady pacemakers. Moreover, the simulation results indicate that the location of nucleation sites may be non-stationary in principle, i.e. these can occur in any place where local synaptic connections have enough resources to excite a minimal critical number of neurons. At the same time, it is quite clear from visual observations that the location of different nucleation sites is not completely independent. In particular, we did not see a tendency to uniform covering the area of the network by randomly arising nucleation sites with increasing the simulation time, although this issue requires further study (e.g., determining the minimal size of a nucleation site and the minimal distance at which two closely located nucleation sites can be distinguished from each other).

In addition, compared to the deterministic case of background currents (see Fig. 7 below and Ref. [29]), one can observe that the propagation speed of spiking activity waves from nucleation sites is relatively slow, as if long-range connections were disabled, although this was not the case. The explanation is that in the absence of background currents the excitability of neurons by synaptic currents is lowered so that more incoming spikes are necessary to make a neuron fire, meaning that the expanding spatial front of synchronous spiking activates a narrower outer neighboring area. As the connections between neurons are effectively less influential, nucleation sites in the stochastic case are activated more independently from each other than in the deterministic one: in the first case, PSs often occur from a few nucleation sites almost simultaneously (see Figs. 4–6), and, in the latter, PSs typically arise from one of a few ‘primary’ steady nucleation sites (Fig. 7) [29, 37]. Based on the observations, it is natural to assume that (i) there exists a certain set of nucleation sites, which is predetermined by the initial distributions of ‘key’ network parameters (see Discussion for details) and (ii) each nucleation site can be activated only by some number of certain combinations of initial (incoming from the outside and/or local) spikes. Then in the stochastic case, on the one hand, due to the reduced excitability of neurons, one needs more spikes to activate a nucleation site and, on the other hand, due to the independent and equal ability of spontaneous spike generation, the number of neurons capable of activating the nucleation site is much larger than in the deterministic case with a small fraction of steady pacemakers. Because of the latter, the stochastic activation of nucleation sites allows a larger number of them to get revealed, compared to the case of constant background currents. Another direct consequence is that introducing the scatter in the probability of spontaneous spiking, i.e., making neurons non-equal in this regard, would lead to decreasing the number of the ‘activating’ neurons and, accordingly, to a smaller number of activated nucleation sites.

Indeed, increasing the scatter of *p_sp_* values in simulations, there is a tendency to decreasing the number of independent nucleation sites, i.e. more and more PSs arise from the same nucleation sites (cp. in Figs. 4–6). Moreover, a few of the nucleation sites retain their former location with the scatter increase, indicating that higher dispersion of neurons over *p_sp_* values effectively enhances the influence of the network connectome on the spatial map of nucleation sites.

The above assumption is also consistent with the simulation results for spatial networks with the ‘hybrid’ distribution of neuronal excitability, when 80% of neurons are deterministic non-pacemakers, i.e. these have subcritical background currents and *p_sp_* = 0, and 20% of neurons are stochastic quasi-pacemakers having *p_sp_* = 0.0005 and zero background currents (Fig. 8). The results show a substantial fraction of stable nucleation sites in this case.

Finally, for the spatial networks the PS periodicity was as follows: 644 ± 61 ms with CV = 0.09 in the *p_sp_* case (Fig. 4), 200 ± 4 ms with CV = 0.02 in the *I_bg_* case (Fig. 7), and 207 ± 5 ms with CV = 0.02 in the hybrid case (Fig. 8). As before, one of the most interesting results has been revealed in the corresponding neuronal ISI distributions (Fig. 9). In contrast to the metric-free binomial networks (see Fig. 2), the ISI distribution in the *I_bg_* case exhibits a pronounced ‘distant’ peak at ISI = 180 ms that approximately coincides with the peak in the hybrid case. In addition, one can see quasi-periodic structure of the ISI distribution at large ISI values in the *I_bg_* case and, less clearly, in the hybrid one. In turn, the hybrid and *p_sp_* cases for the metric networks are qualitatively the same as those for the metric-free networks. The pronounced distant peak in the ISI distributions occurs approximately at the mean value of PS periodicity (see above), if PSs are almost periodic (CV ≪ 1) and have a non-random internal structure (i.e., the spatiotemporal patterns of neuronal spikes during PSs are not fully random, but rather structured and repeatable). Note that in the *p_sp_* case there is also a distant peak at the mean PS periodicity, but it is very small.

**FIG. 9.**
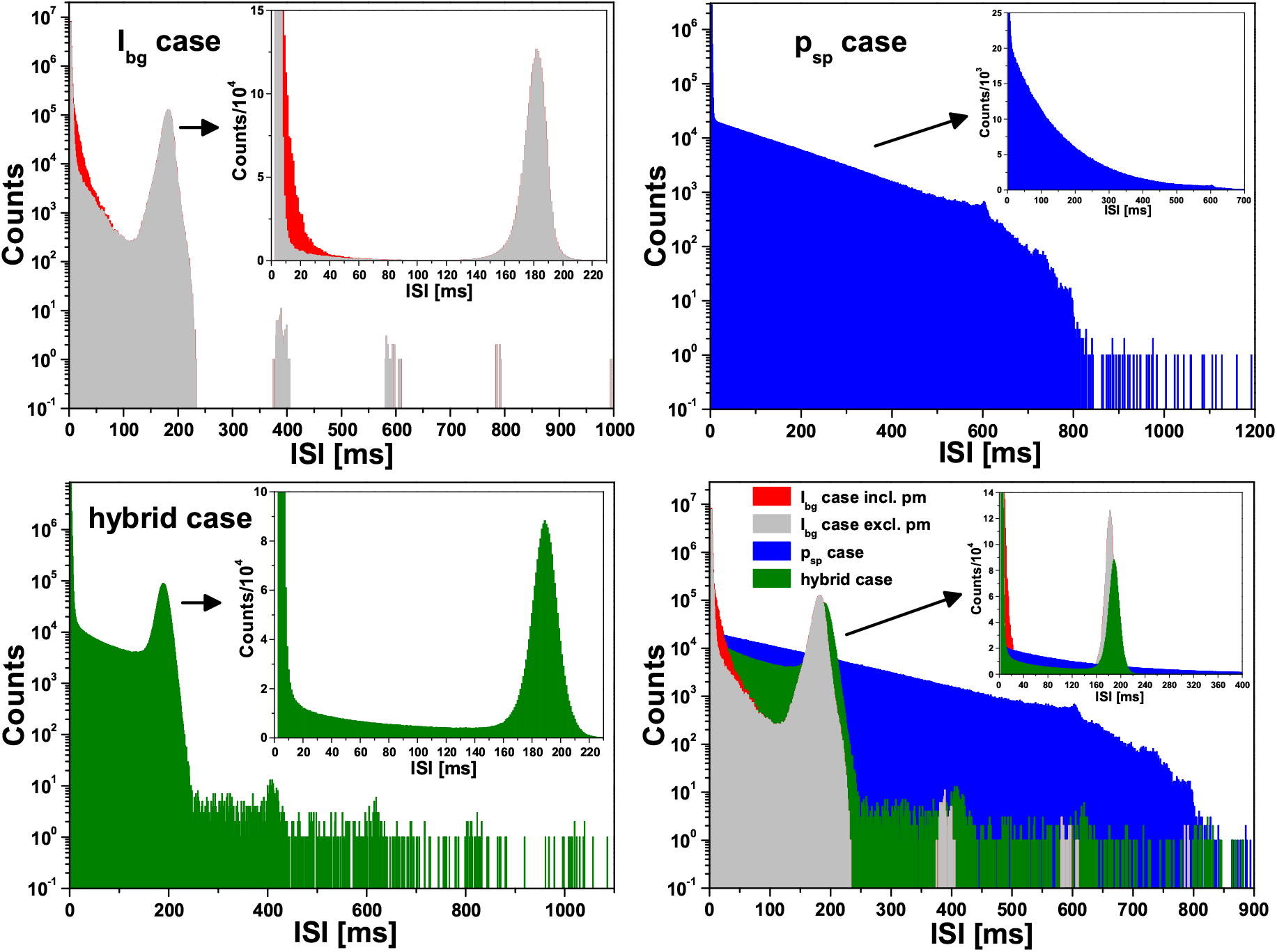
Distributions of Inter-Spike Intervals (ISI) for the ‘spatial’ neuronal networks as those in Figs. 4, 7 and 8. All notations and colors are the same as for Fig. 2. Emphasise that, as in Fig. 2, these distributions are for ‘elementary’ neuronal spikes, not population spikes. One can see that the *p_sp_* case and the hybrid case are qualitatively similar for ‘binomial’ and ‘spatial’ networks. In turn, the *I_bg_* case (i.e. fixed fraction of pacemakers) differs qualitatively: unlike binomial networks, the spatial ones have a large peak at ISI = 180-190 ms that matches the mean timing between successive population spikes, which occur almost periodically (CV ≪ 1, see Fig. 7).

## 3. Discussion

Qualitatively, the repetitive PS regime occurs if (1) the average number of connections per neuron, (2) the average amplitude of the synaptic current pulse, and (3) the probability of spontaneous spike generation (or, in the case of background currents, the fraction of pacemaker neurons) are all greater than certain minimum values. The first condition is for the network connectome, the second is for synaptic interaction, and the third is for the (self-)excitability of network neurons, i.e., for the static variability of internal properties of the neurons. Each pair of the aforementioned minimum values, at the fixed third value, is hyperbolically (i.e. inversely proportional) dependent on each other. Still, the main nontrivial reason for the regime of irregularly repetitive PSs arising in the simulations is that synaptic plasticity nonlinearly modulates the interaction between neurons. Provided this, a dispersion of the three key parameters (in particular, *p_sp_* values) facilitates the occurrence of the PS regime.

The high observed regularity of the PS occurrence in networks of excitatory neurons may lead to a natural association with the respiratory rhythm generation. Indeed, many previous works have shown that this association is quite grounded, by revealing the essential role of emergent network properties for rhythm generation in the pre-Bötzinger complex (PBC) [104–107], which is a relatively autonomous local network of neurons located in the brain stem that produces spiking activity driving the rhythm of mammalian breathing. Unlike the hypothesis on the dominant influence of a fraction of pacemaker neurons (e.g., [108, 109]), with a possible global mutual inhibition of network neurons [110, 111], there are studies related to the PBC and a ‘group pacemaker’ hypothesis that consider only excitatory non-pacemaker neurons with additive noise and dynamic synapses [112–114]. However, these studies use relatively complex biophysical neuron model of Hodgkin-Huxley type, with a rich internal dynamics including limit cycles, which can be activated via stochastic resonance (SR) [115–120]. In comparison with results [112–114], our neuronal network model with stochastic spontaneous activity of neurons is much simpler and, importantly, it shows that nearly periodic PS regime can occur strictly without stochastic resonance. In addition, stochastic spontaneous activity of neurons in the model correlates directly with experimentally-observed stochastic activation of respiratory network activity [121–124].

At the same time, it is worth noting that there exist aperiodic [125–127], self-induced [128, 129], and adaptive [130–132] forms of SR, leading to the non-trivial problem of its identification in neuronal networks. A typical yet insufficient SR signature in this case is the multimodal distribution of neuronal inter-spike intervals (ISI). The fully deterministic version of our network model, with background currents and a fraction of steady pacemaker neurons, indeed exhibits the multimodal ISI distribution at the PS regime, in contrast with its stochastic version. Moreover, multimodality of the ISI distribution is conserved even after excluding the activity of pacemaker neurons from network statistics of neuronal spiking. In turn, the stochastic version of the network model, which is in the focus of this paper, could be related to coherence resonance [133–136] (reviewed in [117]).

It should also be noted that for sufficiently large value of *p_sp_* (such that *ν_sp_τ_ref_* ≳ 1) and sufficiently weak synaptic interaction the identical initial conditions for all neurons and all synapses, together with the same absolute refractory period *τ_ref_* for all neurons, may lead to global long-term transient oscillations of network spiking activity (see Fig. 1 in Ref. [65]) with the period determined by *τ_ref_*. One can imagine that these transient oscillations could lead to sustained ones due to the small but nonlinear influence of synaptic coupling. However, everywhere in this work, we adhere to the condition *ν_sp_τ_ref_* ≪ 1 under which this regime-artifact cannot occur.

Concerning artifacts, it is worth mentioning the reports on noise-induced coherent oscillations in randomly connected neuronal networks [137, 138]: here, ‘noise-induced’ could be understood only in the sense that noise causes dynamics in the system, regardless of any coherent effects (term ‘noise-driven’ seems more appropriate in the context). In turn, the coherent oscillations could occur due to a noise-dependent effect of excessive summation of single contributions (taken in the form of alpha-functions in [138]) to the total incoming synaptic current for each neuron, i.e. the oscillations may be a noise-induced artifact of the model. Despite there exist truly noise-induced collective effects, like SR, we argue that this terminology should be used with caution because of probable deceiving artifacts, which are often unseen in the structure of complex dynamic computational models, particularly the neuronal network models comprising of highly-nonlinear deterministic models of neuronal and synaptic dynamics (e.g., [139]).

Finally, spatial patterns of PS nucleation sites (i.e., coherent spots) in the case of stochastically spontaneously spiking neurons can be naturally attributed to the so-called chimera patterns (or states) [140–144] (reviewed in [145–147]), when two usually mutually-exclusive collective modes coexist simultaneously in different spatial locations.

## 4. Conclusion

We have proposed two (metric-free and metric) neuronal network models that demonstrate the spontaneous occurrence of repetitive population spikes, as well as their unstable nucleation sites in the two-dimensional metric case, despite stochastic spontaneous spiking activity of the network neurons. The results indicate that the regime of nearly-periodic population spikes, which mimics respiratory rhythm, can occur strictly without stochastic resonance and inhibitory interaction between neurons.

## Supporting information

Information about the Supplementary Material

## Data and code availability

The program codes in C for the neuronal network simulator *NeuroSim-TM-2.1* and data processing, the installer for custom-made visualization software *Spatial Activity Monitor*, and all the data shown in this study are assembled into the Supplementary Material available online at: https://doi.org/10.5281/zenodo.4741364

## Author contributions

D.Z. designed and wrote most program codes, performed simulations, analyzed data, prepared graphs, and verified the manuscript. A.P. conceived the study, designed the mathematical model and simulator code, performed simulations, analyzed data, prepared graphs and the supplementary material, and wrote the manuscript.

## Acknowledgments

We acknowledge helpful remarks by two anonymous reviewers. The study was partially funded by the Russian Foundation for Basic Research according to the research project # 17-29-07093. A.P. also thanks the Wellcome Trust for funding his internship at the Neuronal Oscillations Group at the Department of Physiology, Development and Neuroscience at the University of Cambridge in 2014, as the study reported here was initiated during the internship.

---

**Figure S1.**
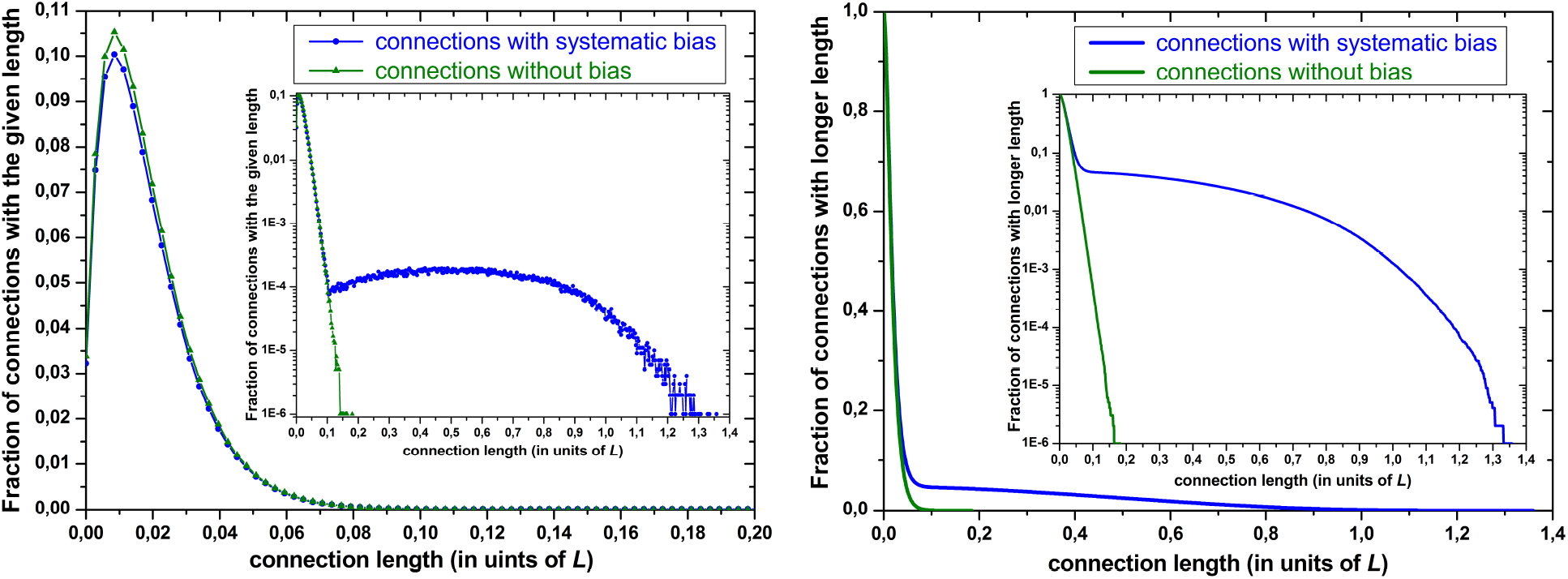
Fraction of connections with the given length (left graph) and with the longer length (right graph): the blue curves represent the data for the network connectome used in simulations for Figures 4–8, with *p_con_*(*r, λ*) = exp(−*r/λ*)+*p*_min_*θ*(*r* −*r*_0_) and *r*_0_ = *λ* ln(1*/p*_min_) (see Eq. (15)), leading to the systematic bias for the fraction of long-range connections. For *p*_min_ = 1*/RAND MAX* ≈ 3 · 10*^−5^* and *λ* = 0.01*L* used in all spatial simulations, one gets *r*_0_ ≈ 0.1*L* that clearly corresponds with what is seen on the graphs. In turn, the green curves are the data for the network connectome used in simulations for Figures S2 and S4 in `\Additional_Simulations` folder, with *p_con_*(*r, λ*) = exp(−*r/λ*) (see Eq. (11)) and without the bias. The inset in each graph is the same graph with the logarithmic scale on the vertical axis. The fractions of connections with the longer length shown on the right graph can be accurately described analytically by Eq. (13) for *n_con_*(*l*) (though the integral can be computed only numerically) substituting there the corresponding expressions for *p_con_*(*r, λ*). Please see subfolder `\Fract_long_conn_Eq13_integral_computation_in_MATLAB` in `\Data_for_FigureS1`… for the details.

**Figure S2.**
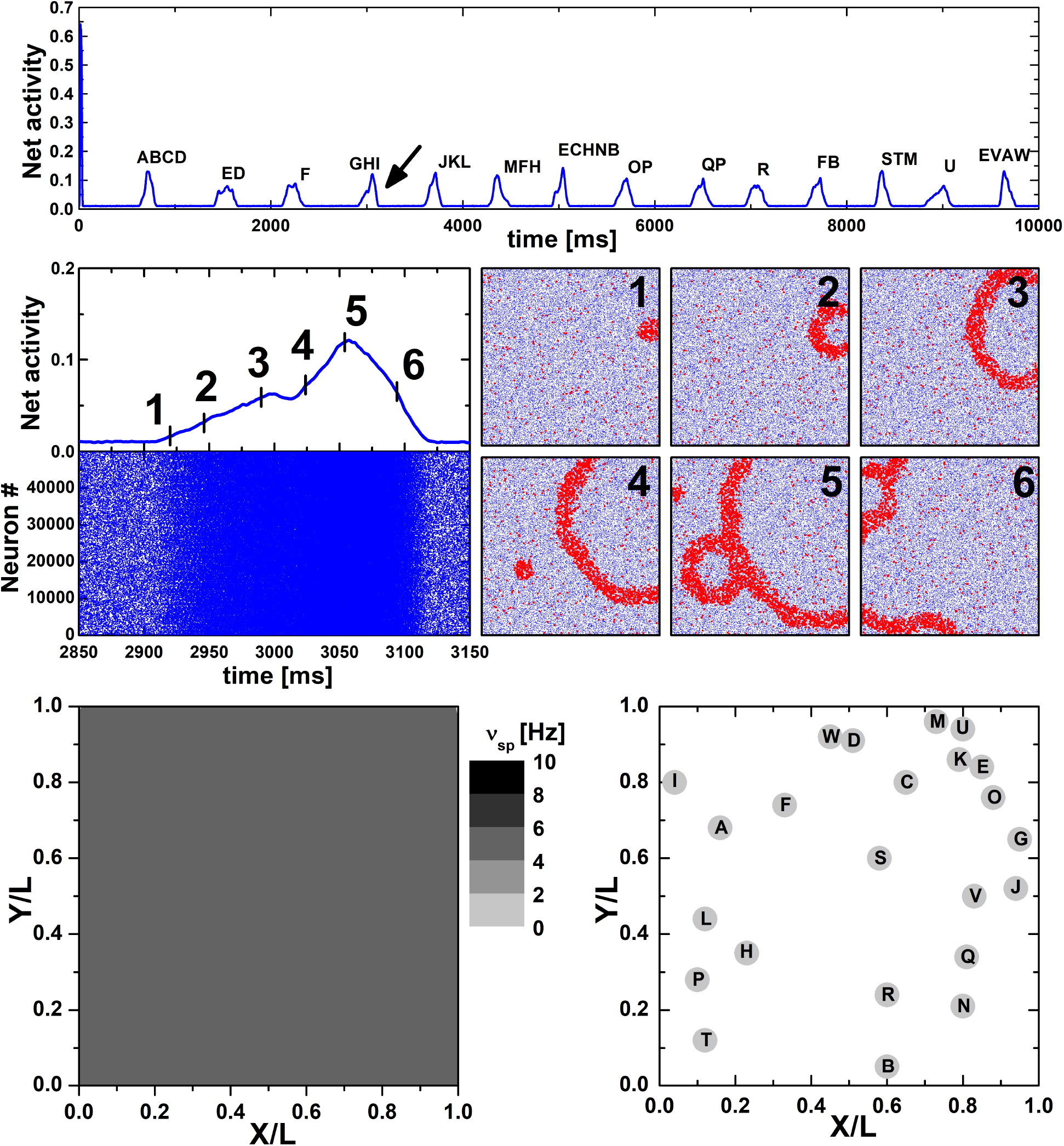
Simulation of spiking activity of a neuronal network consisting of 50 thousand excitatory LIF neurons statistically uniformly distributed over the square *L* × *L*. Spatial coordinates of the neurons are the same as those in Fig. 4 in the main text. Synaptic connections have been formed with probability *p_con_*(*r, λ*) = exp(−*r/λ*), where *λ* = 0.01*L*. This gives 31 ± 6 (mean ± SD) outgoing connections per neuron. Except the network connectome, all other parameters of the simulation are the same as those in Fig. 4. In particular, all neurons have the same value *p_sp_* = 0.0005 of the probability of spontaneous generation of a spike per unit time (Δ*t* = 0.1 ms). **Upper graph:** Network spiking activity, averaged over 2 ms and normalized to the total number of neurons, during 10 seconds of the simulation. Each population spike is denoted by a sequence of Latin letters (uppercase and lowercase letters are not the same), indicating the activation sequence of nucleation sites underlying that population spike. **Middle graph:** LEFT: Network activity (top) and its raster (bottom) during the population spike marked by the arrow in the upper graph. RIGHT: Six snapshots of the instantaneous spatial spiking activity of neurons for the corresponding moments (labeled by the numbers from 1 to 6) of the population spike. Each frame corresponds to the whole area *L* × *L*. Blue dots depict inactive neurons and red dots highlight active neurons. **Bottom graph:** LEFT: Spatial distribution of neurons with a certain value of spontaneous spiking rate *ν_sp_* ≈ *p_sp_/*Δ*t*. As for this simulation all neurons have the same *p_sp_*, the distribution is absolutely uniform. RIGHT: Schematic reconstruction of the spatial pattern of emergent nucleation sites (depicted by filled gray circles) for all population spikes shown in the upper graph.

**Figure S3.**
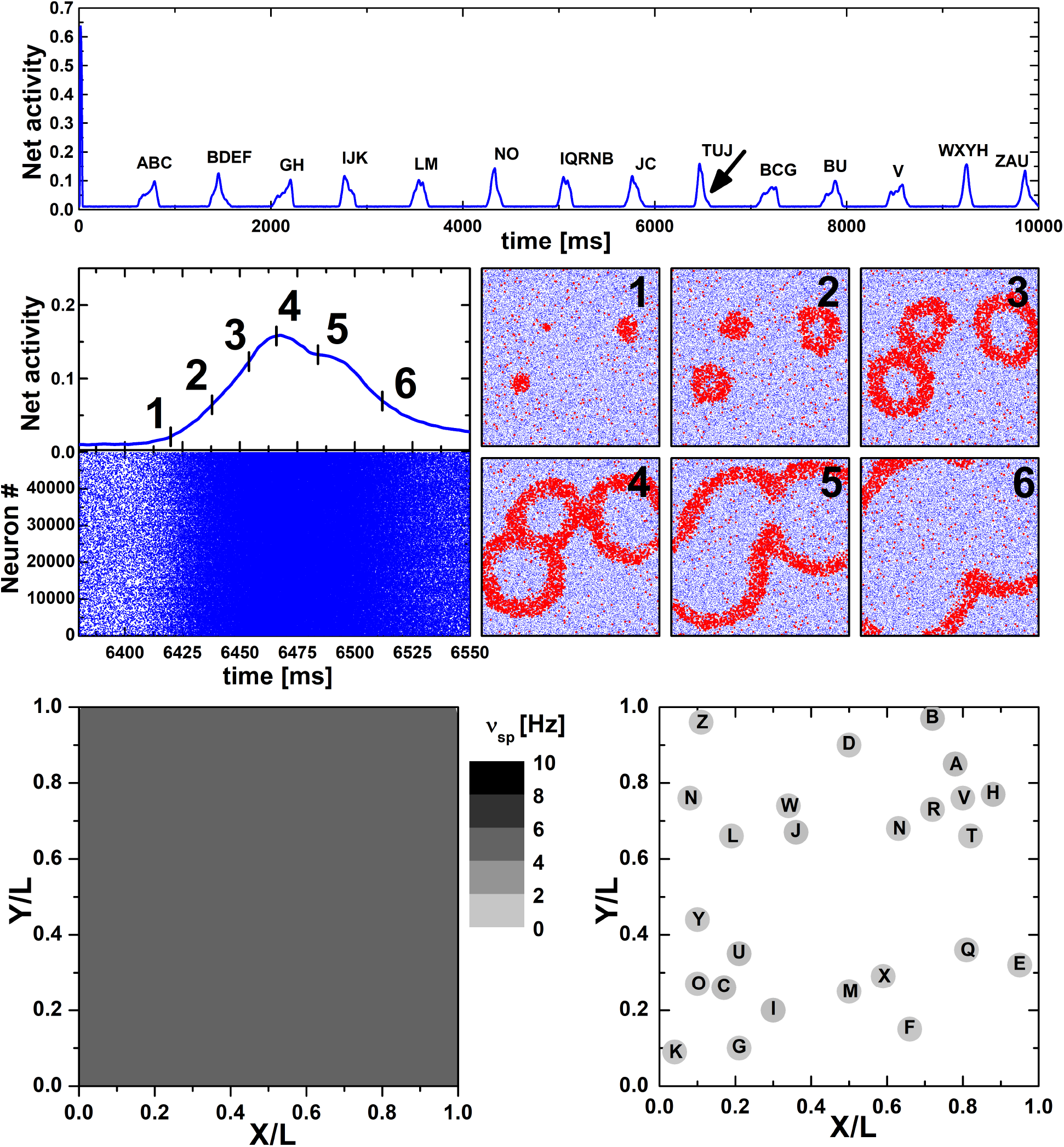
Simulation of spiking activity of a neuronal network consisting of 50 thousand excitatory LIF neurons statistically uniformly distributed over the square *L* × *L*. Spatial coordinates of the neurons are the same as those in Fig. 4 in the main text and in Fig. S2 in the Suppl. Material. Synaptic connections have been formed with probability *p_con_*(*r, λ*) = exp(−*r/λ*), where *λ* = 0.01*L*. Except a newly-generated network connectome, all other parameters of the simulation are the same as those in Fig. 4 and Fig. S2. **Upper graph:** Network spiking activity, averaged over 2 ms and normalized to the total number of neurons, during 10 seconds of the simulation. Each population spike is denoted by a sequence of Latin letters (uppercase and lowercase letters are not the same), indicating the activation sequence of nucleation sites underlying that population spike. **Middle graph:** LEFT: Network activity (top) and its raster (bottom) during the population spike marked by the arrow in the upper graph. RIGHT: Six snapshots of the instantaneous spatial spiking activity of neurons for the corresponding moments (labeled by the numbers from 1 to 6) of the population spike. Each frame corresponds to the whole area *L* × *L*. Blue dots depict inactive neurons and red dots highlight active neurons. **Bottom graph:** LEFT: Spatial distribution of neurons with a certain value of spontaneous spiking rate *ν_sp_*. As for this simulation all neurons have the same *ν_sp_* ≈ 5 Hz, the distribution is absolutely uniform. RIGHT: Schematic reconstruction of the spatial pattern of emergent nucleation sites (depicted by filled gray circles) for all population spikes shown in the upper graph.

**Figure S4.**
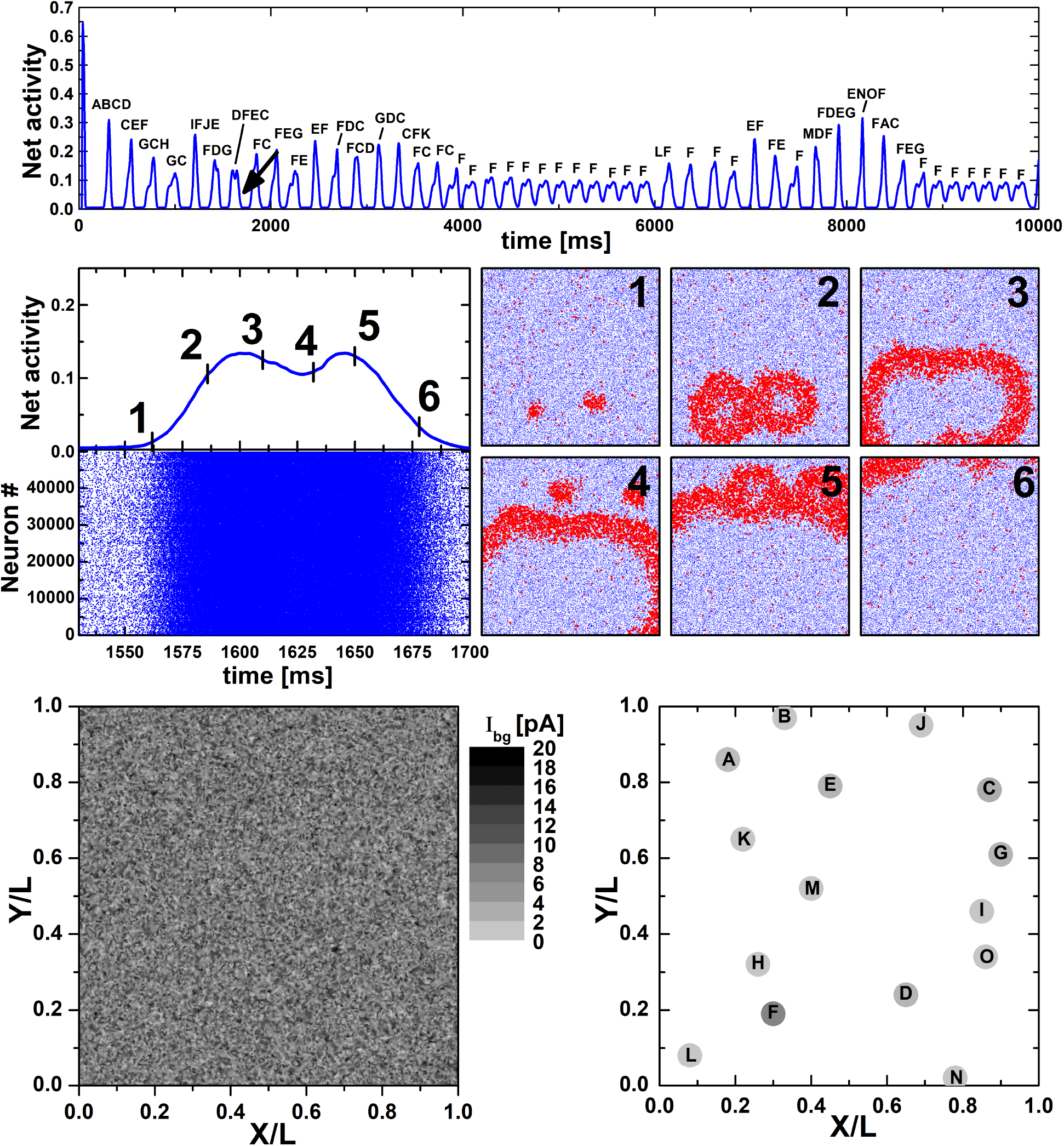
Simulation of spiking activity for the same metric neuronal network as in Fig. S2 except that instead of stochastic spontaneous spiking with probability *p_sp_* all neurons have normally distributed values of the background current *I_bg_* (as in Fig. 1A and Fig. 7 in the main text), resulting in some fraction of steady pacemakers (3.4% of all neurons, see Sect. 2.1). All graphs have the same meaning as those in Fig. 7. **Upper graph:** Network spiking activity, averaged over 2 ms and normalized to the total number of neurons, during 10 seconds of the simulation. Each population spike is denoted by a sequence of Latin letters (uppercase and lowercase letters are not the same), indicating the activation sequence of nucleation sites underlying that population spike. **Middle graph:** LEFT: Network activity (top) and its raster (bottom) during the population spike marked by the arrow in the upper graph. RIGHT: Six snapshots of the instantaneous spatial spiking activity of neurons for the corresponding moments (labeled by the numbers from 1 to 6) of the population spike. Each frame corresponds to the whole area *L* × *L*. Blue dots depict inactive neurons and red dots highlight active neurons. **Bottom graph:** LEFT: Spatial distribution of neurons with a certain value of *I_bg_*. The pacemaker neurons have *I_bg_ > I_c_* = 15 pA. RIGHT: Schematic reconstruction of the steady spatial pattern of emergent nucleation sites (depicted by filled gray circles with the color depth reflecting their relative activation rate) for all population spikes shown in the upper graph.

**Figure S5.**
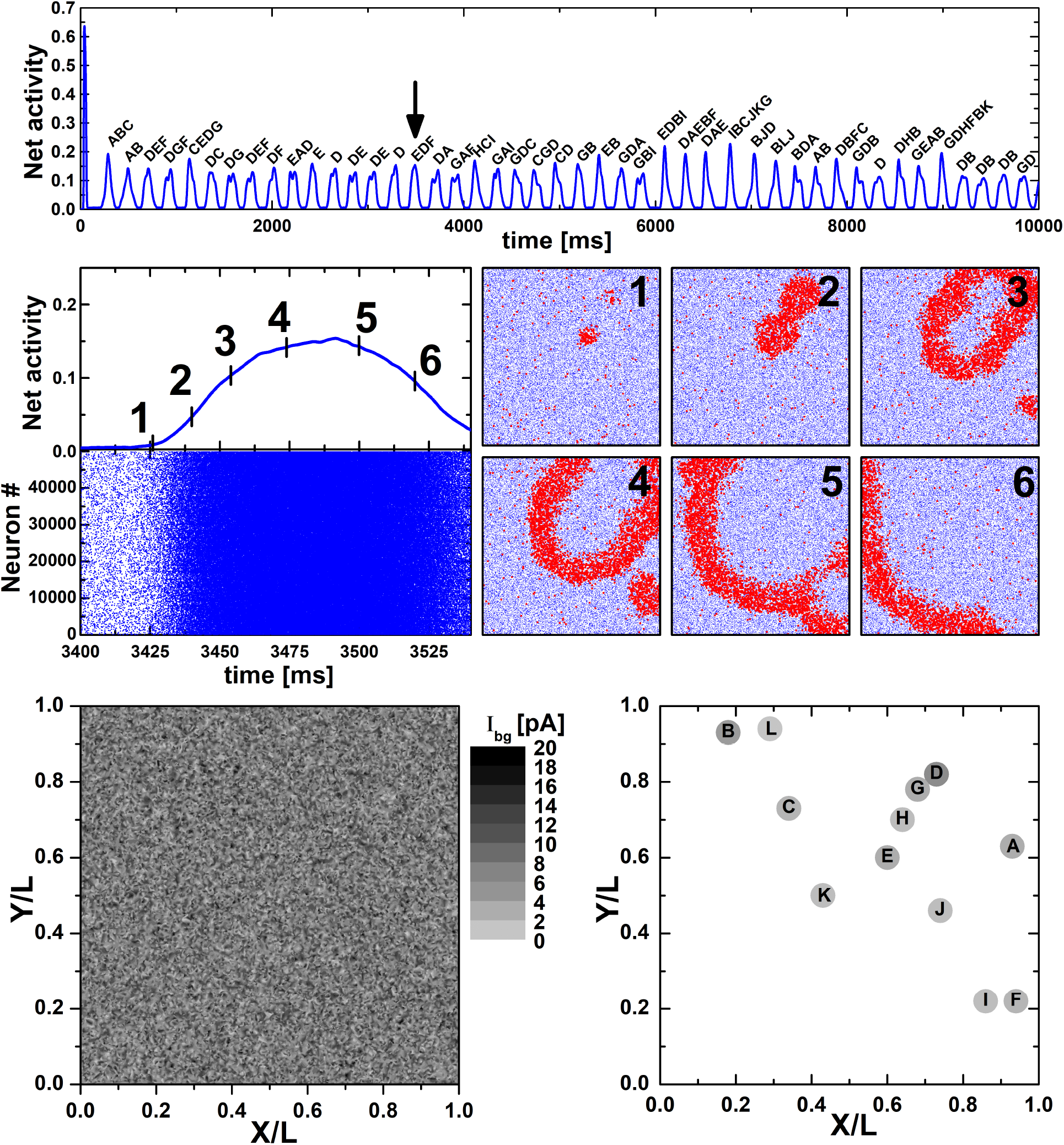
Simulation of spiking activity for the same metric neuronal network as in Fig. S3 except that instead of stochastic spontaneous spiking with probability *p_sp_* all neurons have normally distributed values of the background current *I_bg_* (as in Fig. 1A and Fig. 7 in the main text, and in Fig. S4 in the Suppl. Material), resulting in some fraction of steady pacemakers (3.4% of all neurons, see Sect. 2.1). All graphs have the same meaning as those in Fig. 7 and Fig. S4. **Upper graph:** Network spiking activity, averaged over 2 ms and normalized to the total number of neurons, during 10 seconds of the simulation. Each population spike is denoted by a sequence of Latin letters (uppercase and lowercase letters are not the same), indicating the activation sequence of nucleation sites underlying that population spike. **Middle graph:** LEFT: Network activity (top) and its raster (bottom) during the population spike marked by the arrow in the upper graph. RIGHT: Six snapshots of the instantaneous spatial spiking activity of neurons for the corresponding moments (labeled by the numbers from 1 to 6) of the population spike. Each frame corresponds to the whole area *L* × *L*. Blue dots depict inactive neurons and red dots highlight active neurons. **Bottom graph:** LEFT: Spatial distribution of neurons with a certain value of *I_bg_*. The pacemaker neurons have *I_bg_ > I_c_* = 15 pA. RIGHT: Schematic reconstruction of the steady spatial pattern of emergent nucleation sites (depicted by filled gray circles with the color depth reflecting their relative activation rate) for all population spikes shown in the upper graph.

