## Supplementary material for "Emergent population activity in metric-free and metric networks of neurons with stochastic spontaneous spikes and dynamic synapses": Information about the Supplementary Material

The Supplementary Material (3.8 GB) is available online at: https://doi.org/10.5281/zenodo.4741364

It contains complete data for the Figures shown in the article, including five supplementary Figures in PDF format (see \Additional_Simulations folder). It also contains program codes for the neuronal network simulator *NeuroSim-TM-2.1* and data processing, a brief user guide for the simulator, and the installer for custom-made visualization software *Spatial Activity Monitor*.

For convenience, the supplementary Figures have been included in the preprint after the list of references.
